# Cytomotive actins and tubulins share a polymerisation switch mechanism conferring robust dynamics

**DOI:** 10.1101/2022.09.08.507146

**Authors:** James Mark Wagstaff, Vicente José Planelles-Herrero, Grigory Sharov, Aisha Alnami, Frank Kozielski, Emmanuel Derivery, Jan Löwe

## Abstract

Protein filaments are used in myriads of ways to organise other molecules in space and time within cells. Some filament-forming proteins couple the hydrolysis of nucleotides to their polymerisation cycle, thus powering the directed movement of other molecules. These filaments are termed cytomotive. Only members of the actin and tubulin protein superfamilies are known to form cytomotive filaments. We sought to examine the basis of cytomotivity via structural studies of the polymerisation cycles of actin and tubulin homologues from across the tree of life. We analysed published data and performed new structural experiments designed to disentangle functional components of these complex filament systems. In sum, our analysis demonstrates the existence of shared subunit polymerisation switches amongst both cytomotive actins and tubulins, i.e. the conformation of subunits switches upon assembly into filaments. Such cytomotive switches explain filament robustness, by enabling the coupling of kinetic and structural polarities required for useful cytomotive behaviours, and by ensuring that single cytomotive filaments do not fall apart.

## Introduction

Protein filaments are employed widely in fundamental cell biological processes across the tree of life. Through polymerisation, nanometre-sized protein subunits form larger structures used to organise other molecules at a wide range of scales, up to that of eukaryotic cells many microns across (and sometimes much larger). Polymerisation is therefore one mechanism by which microscopically-encoded information in the genome is able to manipulate the macroscopic environment. Many filament-forming proteins require filament dynamics to function, characteristically using the energy released by inbuilt nucleotide hydrolysis to perform work by pushing and pulling other molecules around. These dynamic filaments have been termed *cytomotive*, to distinguish them amongst the wider class of *cytoskeletal* filaments that perform organising functions (itself a subset of all filament-forming proteins) (Figure 1a) (Löwe and Amos 2009). So far, we know of only two protein families that form cytomotive filaments – the actin and tubulin superfamilies (Figure 1b). These superfamilies encompass the diverse array of homologues of eukaryotic actin and tubulin, found in almost all bacterial and archaeal cells, performing a wide variety of roles in cellular processes (J. Wagstaff and Löwe 2018).

**Figure 1.**
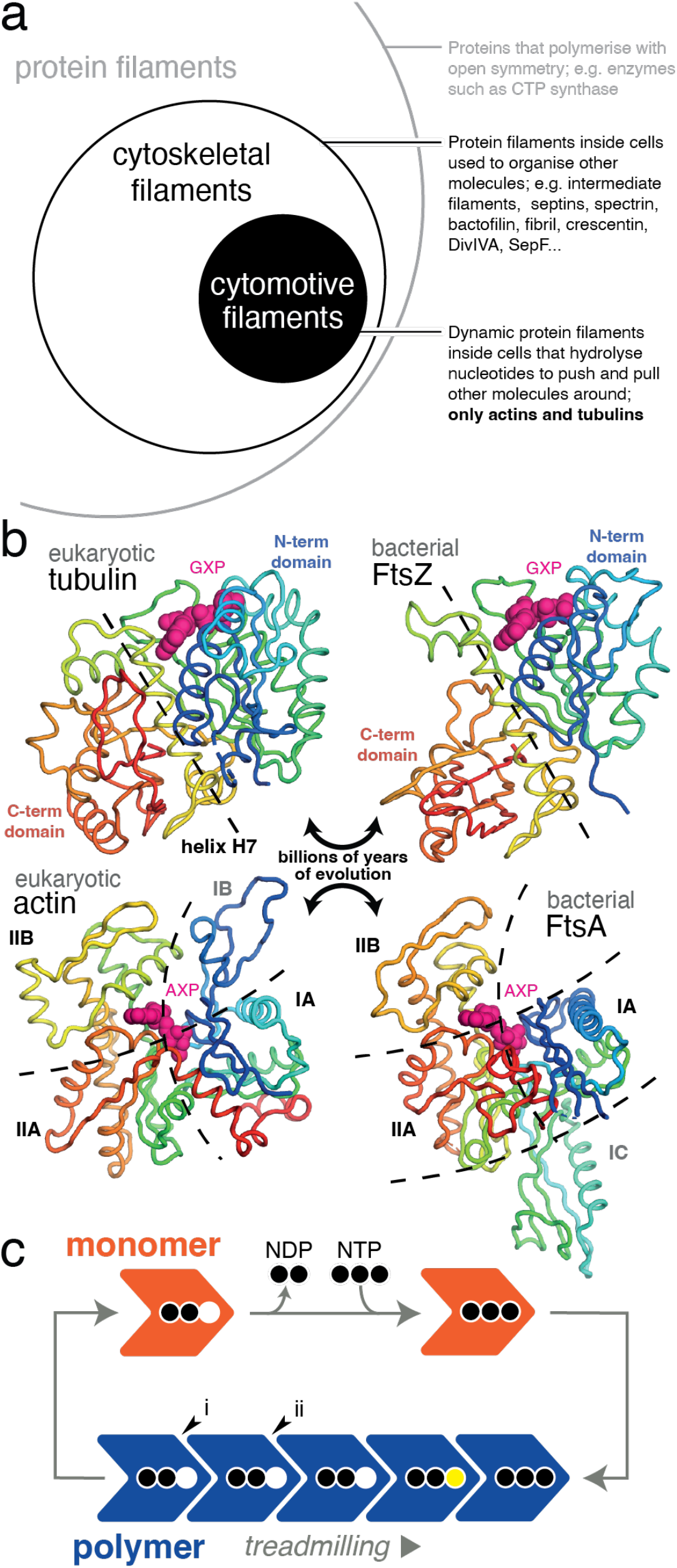
Actin and tubulin superfamilies form cytomotive filaments. (a) Cytomotive filaments are a subset of cytoskeletal filaments, themselves a subset of all protein filaments. Only actin and tubulin superfamily proteins are known to form cytomotive filaments (circles not to scale). (b) Despite billions of years of evolution separating subfamily members in both cases, members of both actin and tubulin superfamilies retain highly conserved structural cores. In the case of tubulins, N- and C-terminal domains are linked by a conserved helix, H7, (yellow). In the case of actins, the core comprises domains IA, IIA and IIB. Structures clockwise from top left are PDB IDs 5NQU, 3VOA, 4A2A, 5JLF. (c) Dynamic properties of cytomotive filaments, such as treadmilling arise due to coupling of nucleotide hydrolysis and polymerisation cycles. The simple model shown is pseudo-isodesmic: the same bonds can be formed/broken upon addition/loss of subunits at either end, such that structural and kinetic polarities are not intrinsically coupled (see also Supplementary Figure S12). In addition, severing at interfaces (i, end) and (ii, middle) should be expected to proceed at the same rate.

Our understanding of how actin and tubulin filaments are able to perform useful work revolves around two dynamic filament behaviours, treadmilling and dynamic instability. Basic models for both behaviours envision a nucleotide-state switch: nucleotide tri-phosphate (NTP)-bound subunits polymerise at growing filament ends, hydrolysis occurs while subunits are within filaments, and nucleotide di-phosphate (NDP)-bound subunits leave from shrinking filament ends – since NDP interfaces are thermodynamically less favourable (Figure 1c) (Alberts 2015; Wegner 1976; Erickson and Pantaloni 1981). While intuitively satisfying, the filaments described by this isodesmic model exhibit two key problems. Firstly, when considering a single-stranded protofilament, loss of an NDP-bound end-subunit is thermodynamically identical to breakage of an NDP interface within the filament and so depolymerisation and breakage are expected to occur at the same rate. Secondly, and less straightforwardly, the isodesmic model does not permit for reliable coupling of structural and kinetic polarities. This means that the growing end of a given filament structure will not always be the same one, relative to the intrinsic geometry of the subunit structure.

Both problems are reduced if a multi-stranded filament is formed, comprising multiple protofilaments bound together by lateral interactions (Alberts 2015; Miraldi, Thomas, and Romberg 2008). Severing of multiple protofilaments becomes thermodynamically different to losing a single end-subunit, and simultaneous formation of different combinations of lateral and longitudinal interactions would appear to offer a way to robustly couple structural and kinetic polarities. Not all cytomotive actins and tubulins are multi-stranded, however (J. Wagstaff and Löwe 2018).

Modelling of eukaryotic actin and microtubule filament dynamics across a range of scales has attracted much attention and has generated insights into many cytoskeleton-associated processes (Wegner 1976; Mitchison and Kirschner 1984; Erickson and Pantaloni 1981; Castle and Odde 2013; Igaev and Grubmüller 2020). Hitherto, modelling has largely been restricted to the well-conserved eukaryotic actin and tubulin proteins and the multi-stranded filaments they form. We posit that this approach has limited progress in understanding the basis of cytomotivity.

Recent years have shown us that actin and tubulin superfamily members from bacteria and archaea also exhibit cytomotive functionality, and that the filaments they form encompass a great deal of structural diversity (J. Wagstaff and Löwe 2018). To improve our understanding of the basis of cytomotivity, we have undertaken a comparative structural biology approach: both assembling and analysing existing data and collecting new structural data.

## Results

### Cytomotive actins share a conformation switch upon polymerisation

Structures have been determined for a number of actin superfamily members apparently representing complete sets of snapshots for their assembly/polymerisation cycles. This includes eukaryotic actin, archaeal crenactin, and bacterial proteins MamK, ParM, FtsA and MreB. All of these proteins share a core of structurally conserved sub-domains (Figure 1b, 2a). The core polymerises in various ways to form diverse filament architectures (Szewczak-Harris and Löwe 2018; J. Wagstaff and Löwe 2018).

**Figure 2.**
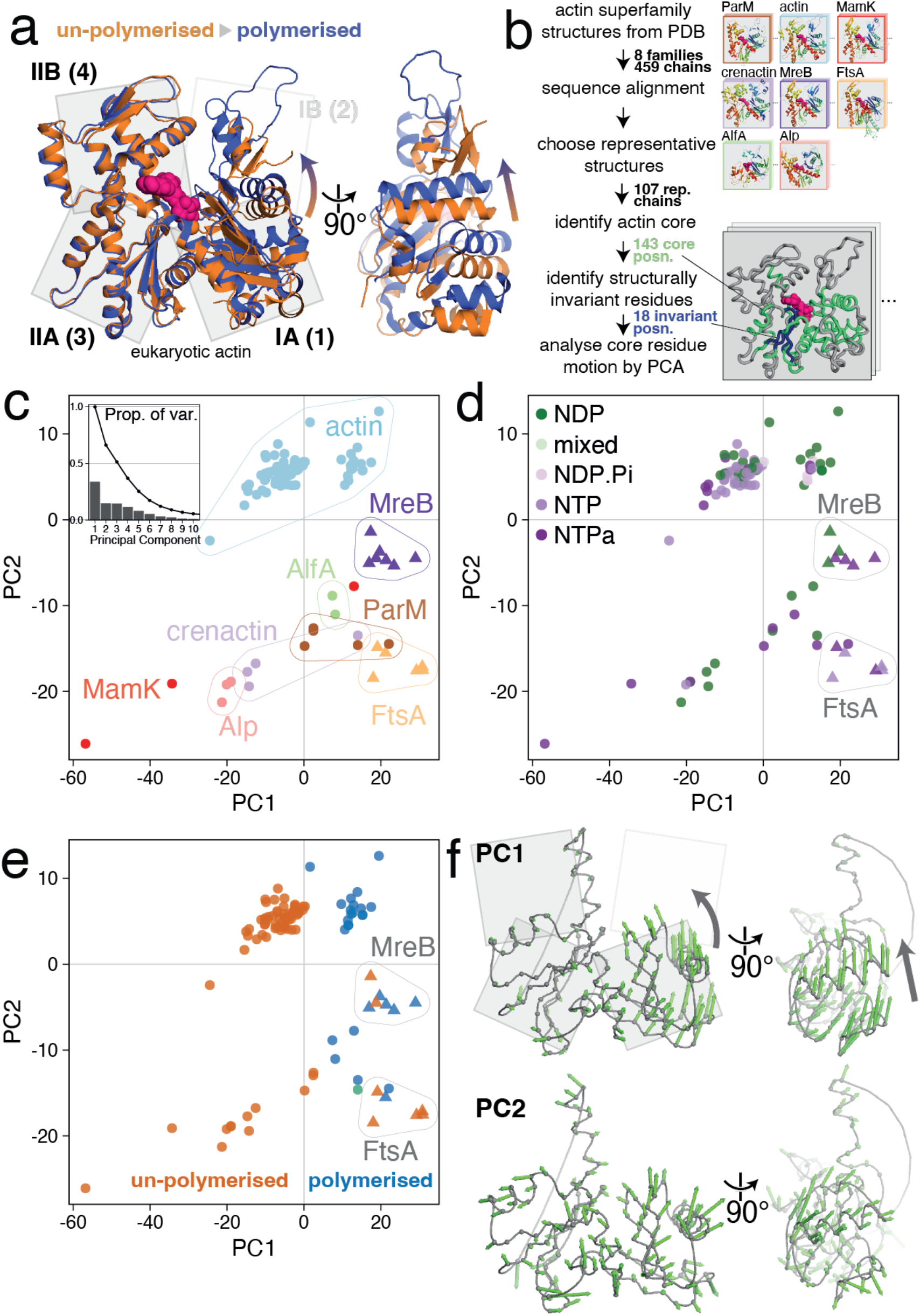
Conformational analysis of actin superfamily structures reveals a conserved subunit switch upon polymerisation. (a) Inspection of eukaryotic actin subunit structures in un-polymerised (PDB 3EL2, orange) and polymerised (5JLF, blue) states reveals the previously characterised actin “propeller twist” conformational change. (b) Pipeline for principal component analysis (PCA) of conformational changes across the actin superfamily. Lower right inset: example structure with identified actin core positions (green), and structurally invariant core (blue). (c) Results of PCA. Representative structures (coloured by subfamily) are plotted in PC1-PC2 subspace. PC2 mostly describes the differences between subfamilies, with further contributions from PC3 (Figure S1). Inset: proportion of variance explained by each component. (d) Identical to (c), but structures are coloured by the hydrolysis state of the bound nucleotide (NTPa = less/non-hydrolysable nucleotide triphosphate analogue). (e) Identical to (c/d), but structures are coloured by polymerisation state (un-polymerised in orange, polymerised in blue; green: PDB 4A62, ParM:ParR, discussed in Supplementary Text S1). PC1 mostly describes the polymerisation state of subunits, with exception of the MreB and FtsA subfamilies (triangles), which form non-cytomotive filaments. (f) Per-position PC loading vectors for PC1 and PC2 are visualised on a representative actin core. Both views as in (a).

To our knowledge, this wealth of structural information has not previously been analysed systematically. To do so, we performed a structure-guided sequence alignment and structural superposition of all Protein Data Bank (PDB) deposited actin superfamily structures, before analysing conformational flexibility at conserved amino acid positions using principal component analysis (PCA) (Figure 2b). PCA reduces complex datasets to a small number of maximally descriptive dimensions and is hence a powerful tool for the analysis of protein conformations (Grant et al. 2006).

The first three, most descriptive, principal components (PC1, 2 and 3) of the actin superfamily dataset explain ∼65% of the variance in amino acid alpha carbon (Cα) positions (Figure 2c, inset; Supplementary Figure S2). PC2 and PC3 largely describe the differences between actin subfamilies (Figure 2c, Supplementary Figure S1, interesting outliers are discussed in the Supplementary Text S1).

If common structural mechanisms underpin the functionality of actin superfamily members, we expect equivalent functional states of different family members to be co-located within PC subspaces. Nucleotide hydrolysis states are often ascribed as defining functional states in protein conformational cycles. Examination of actin PC1-PC2 subspaces does not reveal striking evidence of a correlation between backbone conformation and nucleotide hydrolysis state (Figure 2d). In contrast, subunit polymerisation state is clearly associated with the value of PC1 (Figure 2e). Two almost non-overlapping clusters of structures are seen along PC1: these are polymerised actin and actin-like subunits and monomeric ones - i.e. the structures of assembled actin superfamily subunits are systematically, and discretely, different from those of monomeric subunits. We argue that structural variation within the superfamily corresponding to function can be summarised by performance of the conformational subunit switch, upon polymerisation, as described by PC1.

The subunit switch in eukaryotic actin is well characterised as the “propeller twist” or “monomer flattening” of subdomains IA/B versus IIA/B upon polymerisation [reviewed in (Dominguez and Holmes 2011; Oda et al. 2019)]. The striking finding from our analysis is that PC1, the most descriptive component, succinctly describes the conformational switch upon polymerisation across the entire actin superfamily; despite the billions of years of evolution separating the subfamilies and the significant variation in subdomain composition and longitudinal filament architecture amongst them.

We can visualise PC trajectories on structures, as they describe correlated linear displacements of amino acids (Figure 2f). The conserved switch can be seen in the PC1 trajectories as a closing of the two halves of the monomer, via a hinging between subdomains IIA and IA. The PC2 trajectories are much less concerted, mostly describing differences between families in secondary structure packing.

Bacterial proteins MreB and FtsA (shown as triangles in Figure 2) are an important pair of exceptions to this overall pattern. In these cases, both monomeric and polymerised subunits are placed in the region of PC1-PC2 subspace where we find only polymerised subunits from the other families. These protein families therefore appear to have a “jammed” subunit switch. Crucially, these proteins are thought to form antiparallel, *non-cytomotive* filaments – hence, in cells, these polymers most likely play static, scaffolding roles (Hussain et al. 2018; Nierhaus et al. 2021). Therefore, we note that the subunit switch, performed upon polymerisation, is restricted to cytomotive filament-forming actin superfamily members.

### Solution structures of *Drosophila* tubulin heterodimers confirm presence of subunit switch and absence of nucleotide-driven switch

Our observations of the actin superfamily led us to analyse the tubulin superfamily, the other class of cytomotive filaments.

Despite decades of work, uncertainty remains about the overall conformational landscape of eukaryotic tubulin heterodimers, particularly as to the role played by nucleotide hydrolysis state and to the implications of the sparse sampling of crystal forms resolved for un-polymerised heterodimers (Igaev and Grubmüller 2018; Campanacci et al. 2019). Here, using cryo-EM, we solved solution state structures of recombinant *Drosophila melanogaster* tubulin heterodimers, purified in both GDP and GTP-bound states, in addition to a structure of microtubules (MT), polymerised from the same material (Figure 3a-f).

**Figure 3.**
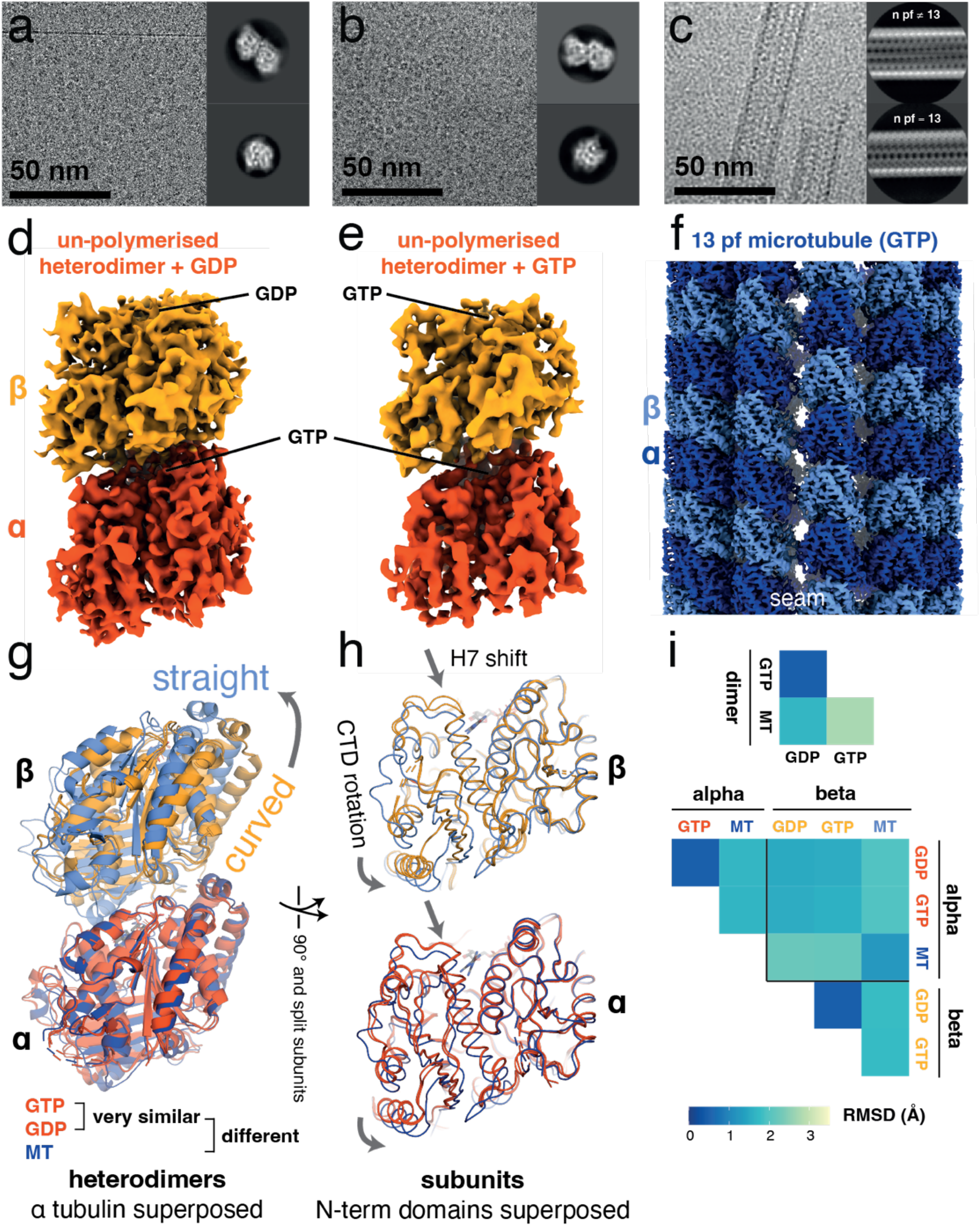
Cryo-EM structures of free/un-complexed *Drosophila melanogaster* (*Dm*) tubulin heterodimers and a microtubule recapitulate classical tubulin assembly switches, and reiterate the absence of a nucleotide state-driven switch. (a) Cryo-EM study of *Dm* tubulin heterodimers prepared with GDP. Representative micrograph and 2D classes. Processing scheme can be found in Supplementary Figure S3. (b) Cryo-EM study of *Dm* tubulin heterodimers prepared with GTP. Representative micrograph and 2D classes. Processing scheme can be found in Supplementary Figure S4. (c) Cryo-EM study of *Dm* microtubules prepared with GTP. Representative micrograph and 2D classes. (d, e) Cryo-EM maps of *Dm* tubulin prepared with GDP or GTP occupying the β-tubulin binding site as indicated. (f) Cryo-EM map of *Dm* tubulin polymerised into 13 protofilament microtubule. (g/h) Comparison of *Dm* models from the dimer maps (orange: dark – α subunit, light – β subunit) and the 13-protofilament MT (blue shades). Structures in (g) are aligned on the N-terminal domain of α-tubulin, structures in (h) are aligned on the N-terminal domains of the respective subunits. (i) Comparison of dimer structures (bottom left) and subunit structures (top right) by Cα RMSD metric, following superposition as in (g) and (h). Un-polymerised heterodimers are very similar to one another.

Structures of GDP and GTP-bound heterodimers (α-tubulin: non-exchangeable GTP, β-tubulin GDP/GTP) were highly similar to each other (Figure 3g-i) and to published crystal structures of other eukaryotic tubulin heterodimers (e.g. all-atom RMSD between GDP-occupied structure by cryo-EM and Darpin-bound heterodimer by X-ray crystallography [PDB 5EYP] is 1.0 Å), confirming that nucleotide hydrolysis state does not play a direct role in determining gross heterodimer conformation (Rice, Montabana, and Agard 2008). Comparison of solution state structures to structures of dimers within MTs recapitulated results seen for other eukaryotic tubulins: a curved-to-straight whole-dimer structural transition upon polymerisation (Figure 3g), and the within-monomer relative rotation of N- and C-terminal domains in both monomers, accompanied by a downward shift of helix 7 (H7) (Fig 3h). The curved-to-straight transition, and the within-monomer between-subunits rotation, have each been proposed as a tubulin assembly switch (Buey, Díaz, and Andreu 2006; Rice, Montabana, and Agard 2008; Michie and Löwe 2006; Ravelli et al. 2004).

The entanglement of the remarkable dynamic properties of MTs, and their structural complexity, being formed of heterodimers, and comprising many protofilaments, presents a formidable barrier to inferring the mechanistic principles underpinning those dynamic properties. We sought to disentangle the mechanisms underlying the idiosyncratic properties of MTs from the more general mechanisms and behaviours of tubulin superfamily protofilaments by examining simpler systems.

### Structures of tubulin superfamily protofilaments support generality of subunit switch

While eukaryotic microtubules are complex in structure and composition, and arose in evolution relatively recently – close in time to eukaryogenesis – the tubulin fold is ancient and can act in isolation as a molecular motor (Nogales et al. 1998). One subunit-thick FtsZ protofilaments, formed of identical monomers, have been observed to treadmill *in vitro* and *in vivo* (Loose and Mitchison 2014; Bisson-Filho et al. 2017; Yang et al. 2017) – and therefore appear to be minimal cytomotive filaments, and a good place to look for clues as to the mechanistic principles underpinning tubulin superfamily cytomotivity.

Many crystal structures of FtsZ have been solved, all of these correspond to monomeric, unassembled FtsZ subunits except for a few crystal forms of *Staphylococcus aureus* (*Sa*) FtsZ (Matsui et al. 2012), in which SaFtsZ crystallises in straight protofilaments extending through the crystals. The “open” conformation of the subunits within the *S. aureus* filaments is markedly different to the “closed” conformation of the monomeric crystal forms from other species. The monomeric closed form is related to the polymeric open form by a CTD rotation, and a shift of H7 – similar to the relationship between corresponding eukaryotic tubulin monomers within curved and straight heterodimers as described in the previous paragraph.

We previously crystallised monomeric *S. aureus* FtsZ in a closed conformation, and produced a low-resolution cryo-EM structure of *E. coli* FtsZ filaments, which showed that subunits within the hydrated filament also adopt the open form seen in *S. aureus* FtsZ crystals (J. M. Wagstaff et al. 2017). Despite extensive efforts we have been unable to produce a high-resolution cryo-EM structure of frozen-hydrated FtsZ filaments. Here, we present a second intermediate resolution structure of a filament from a different organism, *Mycobacterium tuberculosis* (Supplementary Fig S5). This map confirms that polymerised subunits are in the distinctive open conformation. Together, these data demonstrate that single-stranded bacterial FtsZs, much like eukaryotic tubulins, undergo a conformational switch upon polymerisation – an analogy suggested previously (Buey, Díaz, and Andreu 2006), and further supported by the work of others (Y. Chen and Erickson 2011; Fujita et al. 2017).

While the conformational switch upon polymerisation seen in single-stranded FtsZ suggests that multi-strandedness of eukaryotic tubulin microtubules may not be a critical component of the mechanism underlying filament cytomotivity, it does not rule it out.

Straight single protofilaments of eukaryotic tubulin have been observed under certain conditions but their structures have not been resolved at high resolution, e.g. by Elie-Caille *et al*. (Elie-Caille et al. 2007). Similarly, we were unable to generate single-protofilament (1pf) eukaryotic tubulin samples for structural determination.

We thus used a different system of reduced complexity: bacterial tubulin A/B (BtubAB). *Prosthecobacter dejongeii* BtubAB heterodimers form four-stranded (4pf) mini-microtubules, which exhibit dynamic instability and treadmilling (Deng et al. 2017). We designed a mutant of BtubAB, intended to decouple longitudinal polymerisation from lateral assembly interactions by introducing multiple mutations into the M-loop since lateral interactions are exclusively via the “microtubule” M-loop of BtubA. We termed this mutant BtubA*B (Figure 4a). As intended, BtubA*B polymerises in the presence of GTP and GMPCPP to form straight single protofilaments (1pf) (Figure 4a-c, Supplementary Figure S8).

**Figure 4.**
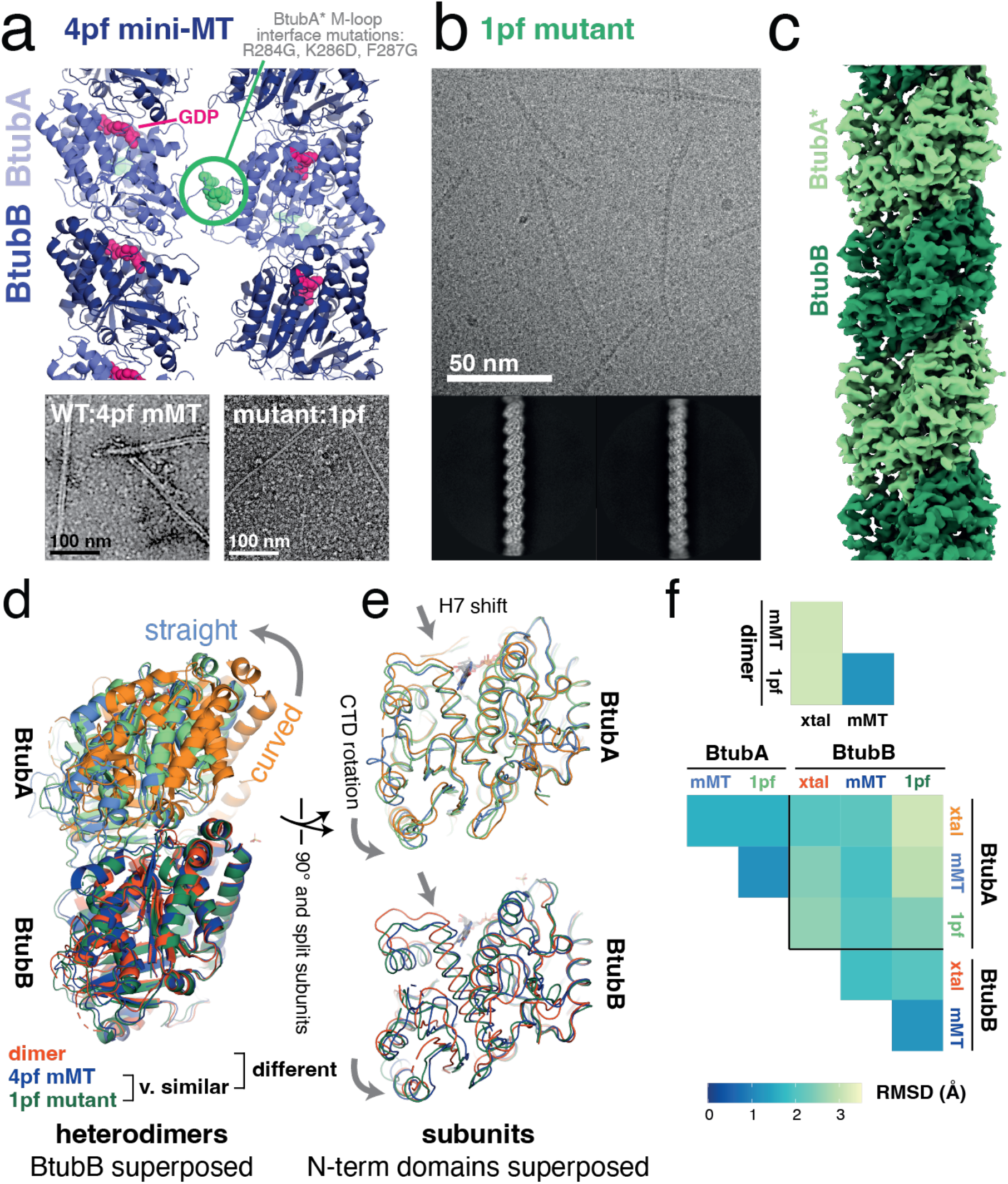
Cryo-EM structures of a single protofilament tubulin reveals a polymerisation-associated conformational switch. (a) M-loop mutations in BtubA prevent lateral interactions between protofilaments (pf). Top: side view of 4-stranded mini-microtubule (mini-MT) structure determined previously by cryo-EM (PDB 5O0C). M-loop residues 284-286 are shown as spheres in green. Bottom: negative stain micrographs: (left) 4pf mini-MT formed by wt BtubAB. (Right) Single protofilament/1pf polymers formed by BtubA*B M-loop mutant (BtubA*: R284G, K286D, F287G). (b) Cryo-EM study of BtubA*B polymerised using GMPCPP. Representative micrograph and 2D class averages are shown. Processing scheme can be found in Supplementary Figure S7. Single filaments are also formed with GTP, see Supplementary Figure S8. (c) Cryo-EM map of a single protofilament formed of BtubA*B polymerised using GMPCPP. (d, e) Comparison of BtubAB models from the dimeric (un-polymerised) crystal form (orange, PDB 2BTQ), the 4pf wt mini-MT (mMT, blue, PDB 5O09) and the single protofilament (1pf, M-loop mutant) solved here (green). Structures in (d) are aligned on the N-terminal domains of BtubB, structures in (e) are aligned on the N-terminal domains of the respective subunits. (f) Comparison of dimer structures (bottom left) and subunit structures (top right) using Cα RMSD metric, following superposition as in (d) and (e). Polymerised heterodimers are highly similar.

Using cryo-EM, we resolved a 3.5 Å reconstruction of single protofilaments of BtubA*B protein polymerised using GMPCPP (Figure 4b, c). Comparison of the 1pf model built into the reconstructed density with the previously determined 4pf mini-MT wildtype (wt) structure (Figure 4d-f) revealed that the 1pf and 4pf polymer structures were essentially identical. Comparison of the polymerised structures and the published crystal structure of wt un-polymerised heterodimers (PDB 2BTQ) as for *Dm* tubulin illustrated the previously characterised curved to straight transition of the dimer upon polymerisation, and the within-subunit inter-domain rotation and H7 shift.

Our explorations of simpler-than-MT tubulin systems therefore appeared to confirm the straightforward conclusion that the conformation changes seen within eukaryotic tubulin heterodimers upon polymerisation are not determined by the multistranded nature of microtubules. Using three different systems also strengthened the case that the within-subunit conformation changes are shared between diverse tubulin superfamily members. To generalise further we proceeded to perform a superfamily structural analysis analogous to that completed for the actins above.

### Subunit switching upon polymerisation is a property of cytomotive tubulins

Tubulin superfamily members are variously employed in a wide range of critical cellular processes across the tree of life (J. Wagstaff and Löwe 2018). Structural studies have been employed to unpick mechanistic details of many these functions, yielding a wealth of data describing the structural plasticity of this ancient fold. As for the actin superfamily, to our knowledge, the tubulin superfamily structural dataset has not previously been systematically analysed in its totality.

All deposited tubulin superfamily structures were therefore analysed in this work, alongside the structures determined here. The methodology described above for the actins was followed: representative structures were selected, residues placed into a common frame of reference via a structure-guided sequence alignment, before determination of a common, invariant structural core and subsequent analysis of conformational differences via principal component analysis (PCA) (Figure 5).

**Figure 5.**
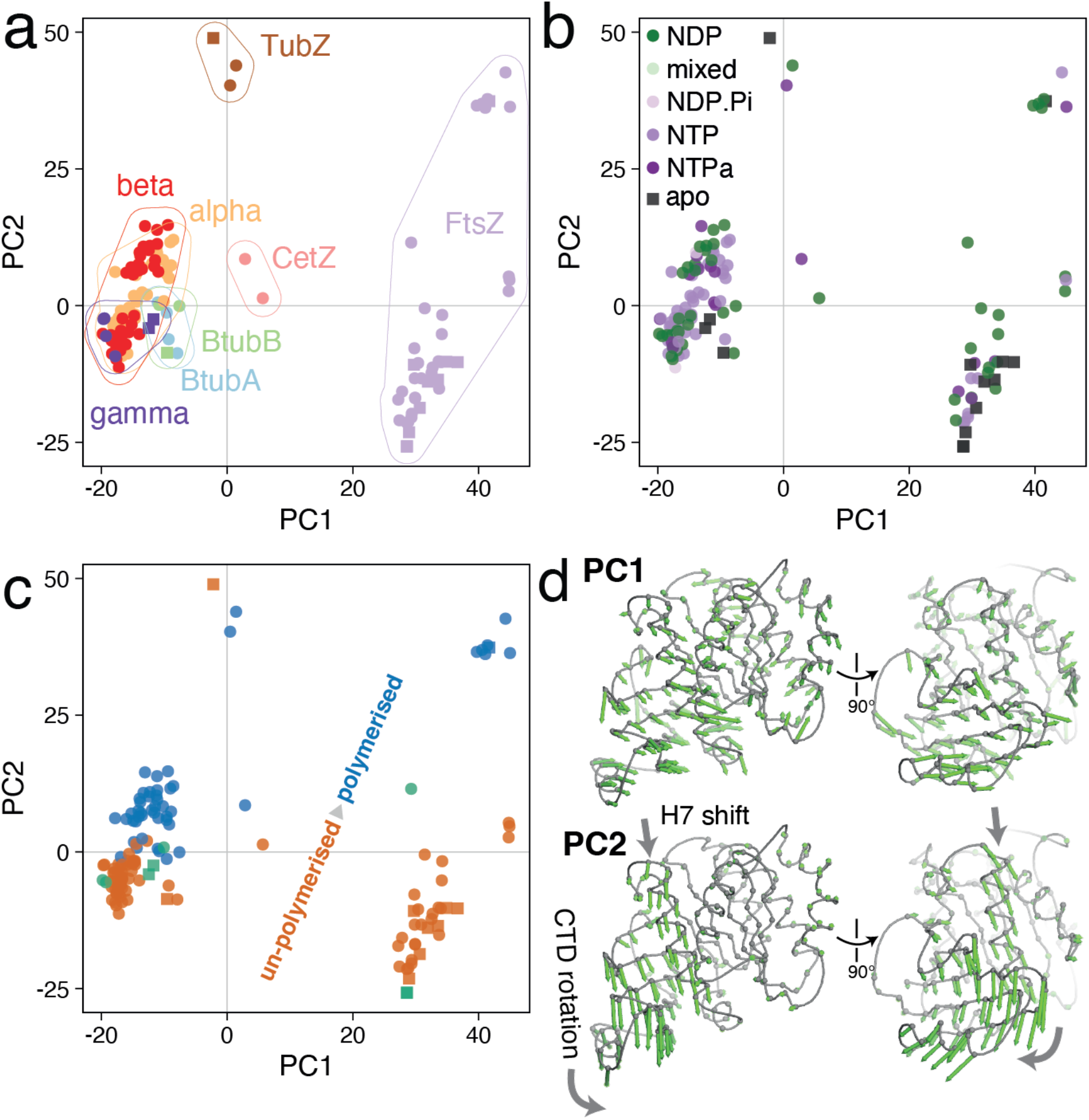
Conformational analysis of tubulin superfamily structures reveals a conserved subunit switch upon assembly. (a) Results of PCA. Representative tubulin structures plotted in PC1-PC2 subspace, coloured by tubulin subfamily. PC1 mostly describes differences between subfamilies. (b) As (a), but structures/points are coloured by nucleotide hydrolysis state (NTPa = non/less-hydrolysable nucleotide triphosphate analogues). (c) As (a/b), but structures/points are coloured by assembly state (un-polymerised – orange, polymerised – blue, “special/ambiguous” – green [more details in Supplementary Text S1]). PC2 mostly describes assembly state. (d) Per position PC loading vectors for PC1 and PC2 are visualised on a representative tubulin core.

As for the actins, the PCA is remarkably successful in compressing the structural variation seen across the superfamily, with PC1 and PC2 alone describing 80% of the variance in Cα positions amongst the representative structures. Examining the PC1-PC2 subspace, it is clear that for the tubulins, PC1 mostly describes differences between subfamilies (Figure 5a), while PC2 describes differences within subfamilies corresponding to conformational changes upon polymerisation (Figure 5b). Again, as for the actins, the hydrolysis state of bound nucleotides does not appear to correlate with position in the PC subspace (Figure 5c) – and is therefore not obviously a determinant of gross conformation.

Plotting of the trajectories corresponding to PC1 and PC2 on representative structures (Figure 5d) shows that PC1 describes the wholesale “widening” of the monomer fold seen between the eukaryotic tubulins and FtsZs, while PC2 indeed describes the within-monomer inter-domain rotation and accompanying shift in H7, observed upon polymerisation of *Dm* tubulin, *Mtb* FtsZ and BtubAB in the structural studies described above.

Unusual structural states can be quickly identified in the PC1-PC2 subspace, details of several of these are discussed in the Supplementary Text S1 and can be inspected in Supplementary Figure S11. Overall, however, the picture is clear: the tubulin superfamily, much like the actin superfamily, appears to possess a shared subunit conformational switch upon polymerisation. The role of conformational changes within the divergent viral TubZ family is less apparent from this analysis, our working hypothesis is that these proteins are (like MreB and FtsA) “jammed” switches, and that their cytomotive properties (Fink and Löwe 2015) are explained by an alternative mechanism driven by the unusual structural interactions between subunits along and possibly across protofilaments via long C-terminal tails.

## Discussion

We set out to improve our understanding of the mechanisms underpinning cytomotive behaviours of protein filaments. Our approach was to complete a structural survey of the actin and tubulin superfamilies, via assembling existing datasets but also by generating new data to fill important gaps. We hypothesised that such an approach would yield insights due to the unusual relevance of structural methods for examining polymerisation as a protein function, and also due to the richness of existing data – with many sets of structures comprising complete, or almost complete, snapshots of functional cycles for individual subfamilies already deposited.

In summary, the approach taken suggested that subunit assembly switches exist within both actin and tubulin superfamilies, in each case the switching mechanism is to a great extent shared amongst the superfamily. In both superfamilies these assembly switches have been identified previously within individual subfamilies – as too in many cases have analogies been drawn between subfamilies (for example in the cases of FtsZ and eukaryotic tubulin (Buey, Díaz, and Andreu 2006), and ParM and eukaryotic actin (Bharat et al. 2015)).

We now present the idea that in both superfamilies the assembly switch is the crucial feature for understanding dynamic filament properties. We ultimately suggest that the cytomotive behaviours common to the two superfamilies are shared *because* of the analogous assembly switches.

Previously, we set out the narrower claim that the treadmilling behaviour of single-stranded polymers of the prokaryotic tubulin FtsZ can be explained by considering the properties of a model filament composed of subunits that perform a conformation switch upon assembly (J. M. Wagstaff et al. 2017). This was subsequently supported by detailed agent-based modelling (Corbin and Erickson 2020). Since then, similar arguments have been advanced for understanding the behaviour of both eukaryotic actin and microtubules (Stewman, Tsui, and Ma 2020; Zsolnay et al. 2020).

The common features amongst these models are summarised in Figure 6. The two challenges outlined above in the Introduction for pseudo-isodesmic models of assembly, namely: filament-splitting occurring at the same rate as end-subunit leaving, and filament kinetic polarity being uncoupled from structural polarity (so that the direction of e.g. treadmilling is defined by the history of the filament, not by the orientation of the subunits relative to the substrate, Supplementary Figure S12) are not faced by models that include a subunit assembly switch.

**Figure 6.**
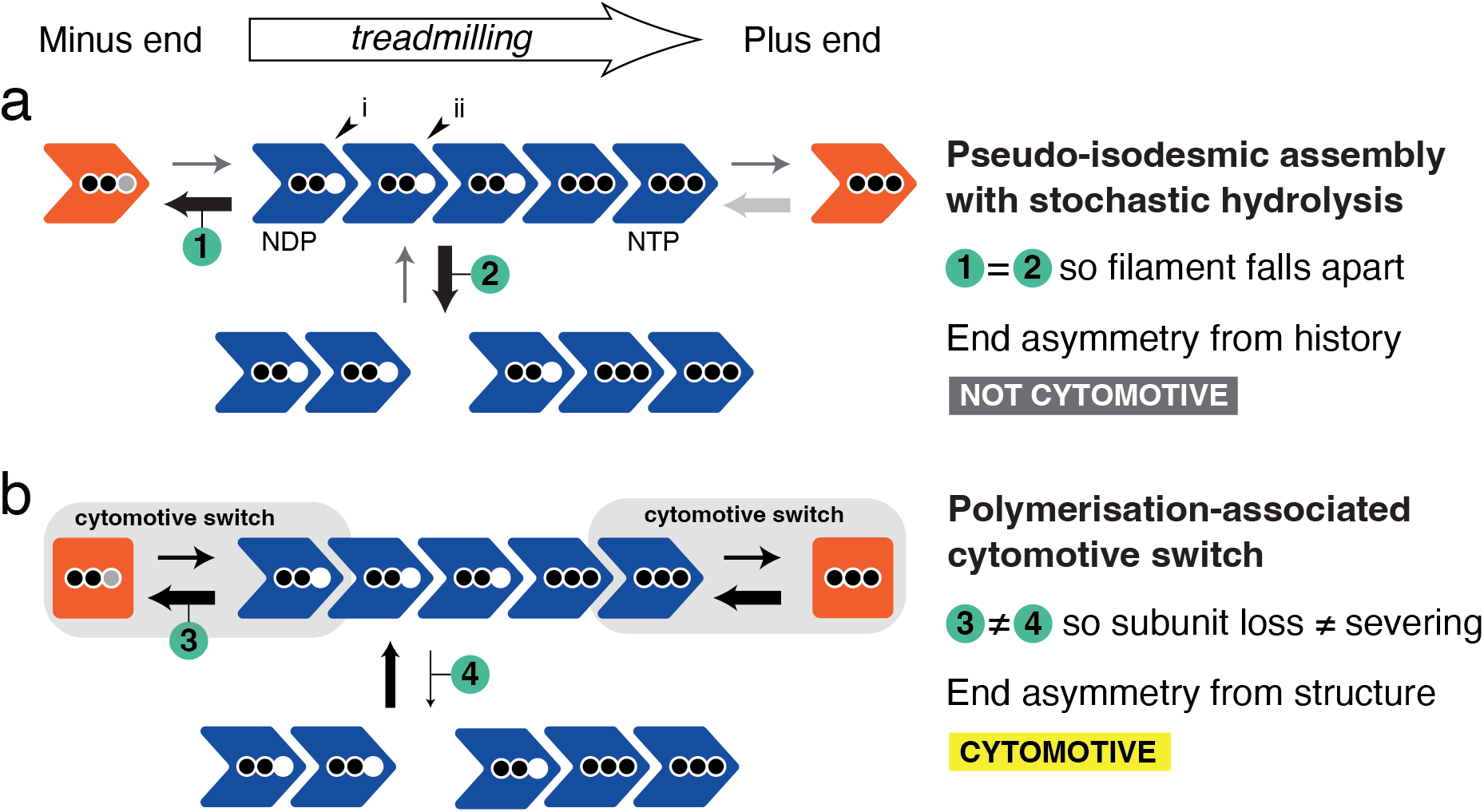
A polymerisation-associated subunit switching mechanism, the “cytomotive switch” is required for robust single-stranded filament dynamics. Two models for protein filament polymerisation are shown. In (a) subunits are rigid. In (b) subunits perform a cytomotive switch, i.e. they have two conformations: one compatible with being in a filament but which is unstable when un-polymerised (blue), and a second that is incompatible with polymerisation but is stable when un-polymerised (orange). Arrows of the same appearance indicate rates that are identical, their widths are proportional to the rates they represent. Two black dots denote NTP-loaded subunits, two black dots mean NDP. A grey dot means exchange of NDP/NTP. The filament in (a) faces two problems, which render it not usefully cytomotive; the filament in (b) employs a cytomotive switch to solve both problems. End asymmetry/coupling of kinetic and structural polarities is explained further in Supplementary Figure S12.

In other words, actins and tubulin proteins have persisted over many billion years during evolution because they are an implementation of a polymerisation-coupled conformation switch that is *required* for them to do useful work in cells that goes beyond a scaffolding “cytoskeleton”. We propose the name “cytomotive switch” to describe polymerisation-associated conformation switches in nucleotide-driven dynamic, cytomotive filaments.

## Acknowledgements

We thank the community for providing the structural data required for our analyses and we apologise that due to space constraints we have not been able to provide citations for structures deposited in the PDB. We thank all members of the LMB electron microscopy facility for excellent EM support, Diamond Light Source for access to and support at the cryo-EM facilities at the UK’s National Electron Bio-imaging Centre (eBIC), funded by the Wellcome Trust, MRC and BBSRC, and T. Darling and J. Grimmett (LMB scientific computing) for computing support. J.M.W. was supported by a PhD scholarship from the Boehringer Ingelheim Fonds. This work was funded by the Medical Research Council (U105184326 to J.L. and MC_UP_1201/13 to E.D.), the Wellcome Trust (202754/Z/16/Z to J.L.), the Human Frontier Science Program (Career Development Award CDA00034/2017 to E.D.). V.J.P-H. is supported by an EMBO postdoctoral fellowship.

## Author Contributions

E.D. and V.J.P-H. purified recombinant tubulin. J.L. purified mutant BtubAB. A.A. and F.K. purified recombinant *M. tuberculosis* FtsZ. V.J.P-H. performed all *Drosophila* tubulin biochemistry. J.M.W. prepared EM grids. G.S. collected one EM dataset. J.M.W. collected all other EM data. J.L. performed EM processing on microtubules. J.M.W. performed all other EM processing. J.L. performed all model-building. J.M.W. performed all sequence/structural analysis. J.M.W. and J.L. conceived of the project. J.M.W. prepared all figures and wrote the manuscript. All authors contributed to editing.

## Methods

All reagents were purchased from Sigma Aldrich unless otherwise specified.

### Protein expression and purification

The amino acid sequences of the proteins used are listed in Supplementary Text S2.

### Recombinant *Drosophila melanogaster* αβ-tubulin heterodimers

A codon-optimised gene for tubulin α1 84B (GenBank entry NM_057424) carrying a tandem N-terminal His_6_-tag and a Protein C epitope tag (PC tag, EDQVDPRLIDGKG) was custom-synthesised (Twist Bioscience) and cloned into a pMT Puro vector (Derivery et al. 2015). The resulting vector was used to generate a D.mel-2 stable cell line adapted to grow in suspension in serum-free Insect-Xpress medium (Lonza). In brief, cells were transfected using Effectene (Qiagen) and selected in Insect-Xpress medium supplemented with 5 μg/ml puromycin (Gibco). After a couple of days of selection, transgene expression was induced by the addition of 0.6 mM CuSO_4_ to the medium. The cells were kept in Insect-Xpress medium supplemented with 5 μg/ml puromycin and 0.6 mM CuSO_4_ for at least a week, keeping the cell density in the 5-25*10^6^ cells/mL range. Cells were collected by centrifugation, washed in 20 mM HEPES pH 7.6, 150 mM KCl, 1 mM CaCl_2_ buffer, frozen in liquid N_2_ and stored at - 80°C. Note that using this procedure only the α-tubulin is overexpressed, and this is enough to recover functional αβ-tubulin heterodimers, as the recombinant α-tubulin associates with endogenous β-tubulin.

To purify the αβ-tubulin heterodimers, the frozen cells were thawed and diluted in tubulin lysis buffer (80 mM K-PIPES, 10 mM CaCl_2_, 10 μM Na-GTP, 10 μM MgCl_2_, 0.12 mg/mL benzamidine, 20 μg/mL chymostatin, 20 μg/mL antipain, 0.5 μg/mL leupeptin, 0.24 mM Pefabloc SC, 0.5 mM PMSF, pH 6.9) and lysed by extrusion using a dounce homogeniser. Lysed cells were rocked for 1 h at 4 °C to ensure complete microtubule depolymerisation, then clarified by centrifugation at 66,000 x *g* for 30 min using a JA 25.50 rotor (Beckman). The clarified lysate was incubated with 2 mL of pre-equilibrated Protein C affinity resin (Roche) for 3 h at 4 °C. The resin was then packed into an empty column (Bio-Rad), washed with 50 mL of tubulin wash buffer (80 mM K-PIPES, 10 μM Na-GTP, 10 μM MgCl_2_, 1 mM CaCl_2_, pH 6.9), 50 mL of tubulin ATP-buffer (wash buffer supplemented with 10 mM MgCl_2_ and 10 mM Na-ATP), 50 mL of low-salt buffer (wash buffer + 50 mM KCl), 50 ml of high-salt buffer (wash buffer + 300 mM KCl), 50 mL of Tween buffer (wash buffer + 0.1% Tween-20 and 10% glycerol) and finally 50 mL of tubulin wash buffer without CaCl_2_. Recombinant tubulin was eluted by incubating the resin with 2 mL of tubulin buffer (80 mM K-PIPES, 10 μM Na-GTP, 10 μM MgCl_2_, 5 mM EGTA, pH 6.9). Multiple elution steps were performed and the fractions were analysed by SDS-PAGE. The protein-containing fractions were pooled and further purified on a Superdex 200 column (GE Healthcare), equilibrated and eluted in tubulin buffer. The final pool was concentrated using AMICON Ultra-4 centrifugal units and used directly for cryo-EM sample preparation. Mass spectrometry analysis revealed an isotopically pure composition of this preparation: α1-84B/ β1-56D (data not shown).

### GDP exchange of *Drosophila melanogaster* αβ-tubulin heterodimers

To exchange the nucleotide present in β-tubulin, concentrated pure αβ-tubulin heterodimers were incubated with 5 mM EDTA for 1 h on ice. After the incubation, 20 mM Na-GDP and 20 mM MgCl_2_ were added to the sample. The exchanged tubulin was then injected onto a Superdex 200 3.2/300 column (GE Healthcare), equilibrated and eluted in 80 mM K-PIPES, 100 μM Na-GDP, 100 μM MgCl_2_, 5 mM EGTA, pH 6.9 to remove any residual GTP present in the sample. The protein was analysed by SDS-PAGE, pooled and concentrated for cryo-EM.

### Mycobacterium tuberculosis FtsZ

For cryo-EM experiments *M. tuberculosis* FtsZ was freshly expressed and purified as previously described, with small modifications (Alnami et al. 2021).

Briefly, full-length FtsZ (coding for residues 1–379), subcloned into expression vector pProEx, was expressed in BL21(DE3) pLysS and cells were harvested by centrifugation. The supernatant was discarded, cells were resuspended in buffer A (50 mM HEPES pH 7.2, 300 mM NaCl, 5% glycerol and 10 mM imidazole) and lysed on ice by sonication. The lysate was centrifuged at 20,000 x *g* for 15 min at 4°C.

The resulting supernatant was loaded into a 5 mL His-Trap FF column, preequilibrated with buffer A. The column was washed with 100 mL buffer B (50 mM HEPES pH 7.2, 150 mM NaCl, 5% glycerol and 50 mM imidazole) and the protein was eluted with buffer C (50 mM HEPES pH 7.2, 150 mM NaCl, 5% glycerol and 250 mM imidazole).

The protein was dialysed overnight at 4°C in the presence of PreScission protease (1 mg of PreScission protease for 50 mg FtsZ), in cleavage buffer (50 mM HEPES pH 7.2, 150 mM NaCl, 1 mM EDTA, 0.01% Tween 20 and 1 mM DTT). Cleaved FtsZ was further purified by applying a second HisTrap FF column and was found in the flow-through. FtsZ was then dialysed in dialysis buffer (25 mM HEPES pH 7.2, 0.1 mM EDTA, 10 mM DDT, 50 mM NaCl and 5% glycerol), concentrated to 15.5 mg/mL, aliquoted, frozen in liquid nitrogen and stored at -80°C.

### Prosthecobacter dejongeii BtubA*B

BtubAB proteins from *Prosthecobacter dejongeii* were co-expressed and co-purified according to previously published protocols (Deng et al. 2017; Schlieper et al. 2005), with some modifications to improve purity given that smaller single protofilaments were to be imaged. The three mutations in the M-loop of the BtubA subunit (R284G, R286D, F287G – the triple mutant is henceforth denoted BtubA*) were designed based on the BtubAB mini microtubule cryo-EM structure [PDB 5O09, (Deng et al. 2017)] to stop interactions with subunits from neighbouring protofilaments. They were introduced by mutagenic PCR (Q5 mutagenesis kit, NEB) of the BtubAB expressing plasmid (Schlieper et al. 2005), using the primers CCGTTGACACCGCCAGACGGCAGTGATGGTGAGGAATTGGGCATTGAG and AGCAAAGGCGCACATGAGGAAGTGCAGCGA, and blunt ligation of the product.

6 L of 2xTY media containing 100 μg/mL ampicillin were inoculated with transformed C41(DE3) *E. coli* cells from three overnight selective TY 90 mm plates. The cultures were grown until mid-log phase at 36 °C and induced with 1 mM isopropyl β-d-thiogalactoside for 3 h at 36 °C, while shaking in 2 L flasks at 190 rpm. The cells were harvested by centrifugation, frozen in liquid nitrogen and stored at -80 °C. The cells were thawed and resuspended in 300 mL of buffer A (20 mM Tris/HCl, 1 mM sodium azide, pH 8.5). Six EDTA-free protease inhibitor tablets (Roche) and small amounts of solid DNase I (Sigma) were added. The cells were opened using a cell disruptor at 25 kPSI (Constant Systems). Cleared lysate was obtained by centrifugation in a Beckman 45 Ti rotor at 35,000 rpm for 30 min. The supernatant was applied to two 5 mL HiTrap Q XL (Cytiva) columns at 5 mL/min, which had been equilibrated in buffer A. Protein elution was achieved using stepwise increases of the concentration of buffer B (buffer A + 1 M NaCl). Most of BtubA*B eluted at 25% buffer B as determined by SDS-PAGE of the resulting fractions. Fractions containing BtubA*B were concentrated using centrifugal concentrators with a 10 kDa molecular weight cut off (Vivaspin 20, Sartorious), and applied to a Sephacryl S300 16/60 column (Cytiva), equilibrated in buffer C (20 mM Tris, 1 mM EDTA, pH 7.5). Fractions were again checked by SDS-PAGE and concentrated as before. The sample was diluted ten-fold with buffer A (pH 8.5) and applied to a Mono Q 4.6/100 column (Cytiva), equilibrated in buffer A. Proteins were eluted with a 40 column volumes linear gradient to 100% buffer B, and most of BtubA*B again eluted at around 25 % buffer B, but with increased purity. Fractions were checked by SDS-PAGE and concentrated as before using centrifugal concentrators to around 500 μL and 42 mg/mL, as determined by the UV absorption of the concentrated sample and a calculated molar extinction coefficient. Aliquots of the sample were flash frozen in liquid nitrogen and stored at -80 °C.

### Cryo-EM sample preparation and imaging

For all imaging, a nominal defocus range of -1.0 - -3.0 μm was used. All Quantifoil/UltrAuFoil grids were purchased from Quantifoil Micro Tools GmbH.

### GTP-bound *Drosophila melanogaster* αβ-tubulin heterodimers

*D. melanogaster* tubulin heterodimers, purified in the presence of GTP as above, were diluted into cold BRB buffer (80 mM PIPES pH 6.9, 1 mM MgCl2, 1 mM EGTA) with GTP (50 μM) to a final concentration of 0.05 mg/mL. 2.5 μL was applied to a gold-on-gold grid (UltrAuFoil R2/2 200 mesh), prepared with a single layer of graphene oxide (Martin et al. 2016). After a wait of 30 s, the grid was blotted on both sides for 4 s and then vitrified by plunge freezing using a Vitrobot Mark IV (FEI) into liquid ethane maintained at 93.0 K using an ethane cryostat (Russo, Scotcher, and Kyte 2016). The Vitrobot chamber temperature was set to 10 °C and humidity to 100%. Micrograph movies were collected using a Titan Krios TEM (FEI) operating at 300 kV, with a K3 detector (Gatan Inc.). Pixel size was 0.86 Å, dose per frame was adjusted to 1 e-/Å^2^, 40 frames were recorded. A total of 4,042 movies were collected over two sessions.

### GDP-bound *Drosophila melanogaster* αβ-tubulin heterodimers

*D. melanogaster* tubulin heterodimers, purified in the presence of GTP and then exchanged into buffer containing GDP as above, were diluted into cold BRB buffer (80 mM PIPES pH 6.9, 1 mM MgCl_2_, 1 mM EGTA) also with GDP (50 μM) to a final concentration of 0.05 mg/mL. Grids were prepared as for the GTP-bound sample. Micrograph movies were collected using a Titan Krios TEM (FEI) operating at 300 kV, with a K3 detector (Gatan Inc.). Pixel size was 0.86 Å, dose per frame was adjusted to 1 e-/Å^2^, 40 frames were recorded. A total of 5,724 movies were collected in one session.

### Drosophila melanogaster microtubules

To prepare dynamic *D. melanogaster* microtubules, 5 μM of purified αβ-tubulin heterodimers were diluted in polymerisation buffer (80 mM K-PIPES, 10% DMSO, 1 mM GTP, 1 mM MgCl_2_, pH 6.9), supplemented with 10 μM Taxol to favour microtubule nucleation. 5 μl of this reaction were then added to a 35 μl solution of 25 μM αβ-tubulin heterodimers in polymerisation buffer (without Taxol) and incubated at 37 °C for 20 min to induce microtubule polymerisation. Note that the final Taxol concentration in this sample is c.1 μM. After the incubation, the protein was layered on top of a cushion solution (80 mM K-PIPES, 1 mM GTP, 1 mM MgCl_2_ and 60% glycerol, pH 6.9) and centrifuged at 100,000 xg for 30 min at 37° C using a warm TLA100 fixed angle rotor (Beckman). After the spin, the top solution was removed and the interface with the cushion solution was washed with EM buffer (80 mM K-PIPES, 1 mM GTP and 1 mM MgCl_2_, pH 6.9). The cushion solution was then removed and the microtubule pellet was washed with 3 × 100 μL warm EM buffer to remove any residual glycerol from the solution. The pellet was then resuspended in warm EM buffer and 3 μL of this sample was applied to a carbon on gold grid (Quantifoil R2/2 Au 200 mesh), recently glow-discharged for 1 min at 30 mA. After a wait of 30 s the grid was blotted on both sides for 4 s and then vitrified by plunge freezing using a Vitrobot Mark IV (FEI) into liquid ethane maintained at 93.0 K using an ethane cryostat (Russo, Scotcher, and Kyte 2016). The Vitrobot chamber temperature was set to 37 °C and humidity to 100%. Micrograph movies were collected using a Titan Krios TEM (FEI) operating at 300 kV, with a K3 detector (Gatan Inc.). Pixel size was 1.08 Å, dose per frame (32 after electron-event representation [EER] fractionation) was adjusted to 1.1 e^-^ /Å^2^. A total of 2,010 movies were collected in one session.

### Mycobacterium tuberculosis FtsZ

*M. tuberculosis* FtsZ, purified and stored as above was thawed on ice before being diluted to a final concentration of 0.2 mg/mL in ice cold buffer HMK100 (Osawa, Anderson, and Erickson 2009) (50 mM HEPES, 100 mM potassium acetate, 5 mM magnesium acetate, 1 mM EGTA, pH 7.7), GMPCPP (Jena Bioscience) was added last to a final concentration of 0.5 mM. The sample was mixed by pipetting, incubated for 2-5 min at 20 °C, and mixed again before 3 μL was applied to a glow discharged (1 min at 40 mA) grid, either carbon on Cu/Rh (Quantifoil R2/2 200 mesh) or gold on gold UltrAuFoil R1.2/1.3 300 mesh. Without wait, the grid was blotted on both sides for 4 s and then vitrified by plunge freezing using a Vitrobot Mark IV (FEI) into liquid ethane maintained at 93.0 K using an ethane cryostat (Russo, Scotcher, and Kyte 2016). The Vitrobot chamber temperature was set to 10 °C and humidity to 100%. Movies with 0° stage tilt were collected using a Titan Krios TEM (FEI) operating at 300 kV, with a K3 detector (Gatan Inc.). Pixel size was 0.86 Å, dose per frame was adjusted to 1 e-/Å^2^, 40 frames were recorded. A total of 13,020 movies were collected in two sessions. Movies with 40° stage tilt were collected using a Titan Krios TEM (FEI) operating at 300 kV, with a K2 detector (Gatan Inc.). Pixel size was 1.47 Å, dose per frame was adjusted to 1 e-/Å^2^, 40 frames were recorded. 660 movies were collected in one session.

### Prosthecobacter dejongeii BtubA*B

Mutant *P. dejongeii* BtubA*B protein purified and stored as above was thawed on ice before diluting to a final concentration of 0.21 mg/mL in ice cold buffer HMK100 (Deng et al. 2017) (50 mM HEPES, 100 mM potassium acetate, 5 mM magnesium acetate, 1 mM EGTA, pH 7.7), with GMPCPP (Jena Bioscience) or GTP added last to final concentrations of 0.5 mM or 5 mM respectively. Samples were mixed by pipetting, incubated at 20 °C for 45 s (GMPCPP) or 2 min (GTP), mixed again and then 3 μL was applied to a freshly glow discharged (1 min, 40 mA) gold on gold grid (UltrAuFoil R1.2/1.3 300 mesh). Without wait, the grid was blotted on both sides for 4 s and then vitrified by plunge freezing using a Vitrobot Mark IV (FEI) into liquid ethane maintained at 93.0 K using an ethane cryostat (Russo, Scotcher, and Kyte 2016). The Vitrobot chamber temperature was set to 10 °C and humidity to 100%. Movies were collected using a Titan Krios TEM (FEI) operating at 300 kV, with a K3 detector (Gatan Inc.). Pixel size was 1.1 Å, dose per frame was adjusted to 1.1 e-/Å^2^, 40 frames were recorded. For the GMPCPP sample, a single grid was imaged. 4728 movies were collected with 0° stage tilt, 1265 were collected with 40° stage tilt. For the GTP sample, 5311 movies were collected on a single grid with 0° stage tilt.

### Cryo-EM data processing

Unless stated otherwise, processing was with Relion 3.1 (Zivanov et al. 2018). CryoSPARC was version 3.1 (Punjani et al. 2017).

### *Drosophila melanogaster* αβ-tubulin heterodimers

#### GTP-bound sample

A schematic processing pipeline can be found in Supplementary Figure S4. Two datasets were collected from a single grid and treated independently until the merging point indicated. Micrograph movies were imported and motion-corrected using the algorithm implemented within Relion. CTF estimation was performed using CTFFIND4 (Rohou and Grigorieff 2015). Micrographs were filtered for high-resolution information content using Thon rings, and for ice thickness by calculating average pixel intensity in a central region, using the mrcfile.py library (Burnley, Palmer, and Winn 2017). Particles were picked using Relion’s Gaussian blob picker, before extraction in boxes of 110^2^ pixels at a nominal 1.72 Å/pixel. Several rounds of 2D classifications were performed, using varying mask diameters and class numbers. Smaller masks were useful for gathering top views of the oblong tubulin heterodimer (for similar approach see (Herzik, Wu, and Lander 2019). Initial 3D refinements used a 20 Å low pass filtered tubulin heterodimer crystal structure (2Q1T). Particles were re-centred/re-extracted in boxes of 148^2^ pixels at a nominal 1.27 Å/pix before 3D refinement and Bayesian polishing, in boxes of 148^2^ pixels using a calibrated pixel size of 1.244 Å/pix. The best particles were recovered by 3D classification without alignment, with varying numbers of classes and values of the regularisation parameter T. Datasets were merged for a final 3D refinement of 34k particles, yielding a 3.5 Å reconstruction.

#### GDP-bound sample

A schematic processing pipeline can be found in Supplementary Figure S3. The overall approach was very similar to that employed for the GDP-bound sample. Refinement of 69k high quality particles yielded a final reconstruction at 3.2 Å.

### Drosophila melanogaster microtubules

Since we reconstructed “naked” microtubules from 2D projection images, lacking any additional subunits indicating the positions of each of the α/β-tubulin heterodimers, we employed an adaptation and extension of the method employed by Lacey et al. (Lacey et al. 2019) to be able to deal with the 26-fold pseudosymmetry of the 13 protofilament microtubules imaged. Two-fold pseudo symmetry arises because of α/β-tubulin being very similar in structure, and additional 13-fold pseudosymmetry arises because the 13 protofilaments are not equivalent since microtubules are not truly helical and have a “seam”, where the B-lattice becomes an A-lattice.

All image processing was done in Relion 3.1 (Zivanov et al. 2018). On import, each electron-event representation (EER) exposure was dose-fractionated into 32 sub-frames that were then aligned against each other to yield motion-corrected images, using 8 and 5 tiles in X and Y. Subsequently, CTF parameters for each image were determined using CTFFIND 4.1 (Rohou and Grigorieff 2015). 1770 images were retained after filtering out those images that produced poor CTF fits as determined by CTFFIND’s resolution estimation. 1222 particles were picked manually and 2D classified after extraction particle images in boxes of 270^2^ pixels and two-fold binning (pixel size 2.16^2^ Å^2^). Six classes were used as the reference for automatic picking of helices as implemented in Relion, and 354,399 particles were picked. Extraction as before and subsequent 2D classification revealed some classes that were not 13 protofilament microtubules, as obvious from by the lack of complete co-linearity of the protofilaments with the MT axis. Only those classes showing fine details and also no twist of the protofilaments were retained, leading to a dataset of 223,884 particle images.

Using these particle images, a fully pseudo-symmetrised reconstruction was calculated with the helical parameters: twist -27.8° (∼360° / 13) and rise 9.51 Å (∼43Å * 3 / 13). This averages all subunits onto all others, disregarding the seam and the differences between α and β-tubulin. This, and all subsequent 3D reconstructions used an overall mask approximately four dimers long and selecting the microtubule wall, only. The resulting reconstruction at 4.4 Å resolution (after masked post processing in Relion) allowed the placing of five tubulin dimers (PDB 3J1T) in one randomly chosen protofilament. From the placed atomic model of a protofilament a 20 Å-filtered map was generated in Chimera (Pettersen et al. 2004), which was used to create a mask that covers a single protofilament. At this point, particle images were re-extracted in boxes of 400^2^ pixels and binned by 2/3 (box 266^2^ pixels, pixel size 1.624^2^ Å^2^). Using the Relion “--local_symmetry” option [see (Lacey et al. 2019) for details], describing the 13-fold protofilament symmetry of MTs, and also overall helical symmetry, now following the heterodimers (twist = 0° and rise = 86.9 Å), a new reconstruction was calculated (resolution 4.0 Å resolution after masked gold standard post processing in Relion) after particle polishing and 3D classifications, based on 44,105 particle images. The overall number of particle images was reduced intently at this step to enable the large symmetry expansion, which followed next.

Using the command “relion_particle_symmetry_expand --helix 1 --i run_data.star --o run_data_expanded26.star --twist 27.692307 --rise -10.0335231 --asu 26” each particle image was added 25 times in all possible other positions within the 26-fold pseudo-redundant MT symmetry. In parallel, tubulin dimers from PDB 3J1T were fitted into the reconstructed map and only atoms from the loop filling the pocket in β-tubulin (residues 359-372) equivalent to the Taxol binding pocket in α-tubulin were left. This atomic model of the most significant difference between α- and β-tubulin was used to subtract map densities in all particle images (which were 26-fold symmetry expanded), that are not in or close to the Taxol binding pocket in α-tubulin or the equivalent pocket in β-tubulin. Several rounds of 3D classification without alignment were then run on these subtracted particle images, until the MT symmetry emerged in the best classes, which meant that only every second pocket along each protofilament was occupied by density (representing residues 359-372 in β-tubulin) and that the seam, where the pattern of alternating pockets changes, was clearly identifiable. Not allowing alignment was the most important setting here: no rotational/translational alignment of particles, only moving particles between classes, means the particles from the symmetry expansion were simply sorted into the correct class that has the right positioning of the seam and along the protofilaments since the symmetry expansion provided all possible versions of each particle image in the dataset. Note that “–local_symmetry” was used in these classification runs since it significantly enhanced the signal by averaging correctly over all subunits. The final map was calculated from 39,594 particle images and was not using “gold standard” separation of half datasets, for purely technical reasons of Relion’s implementation. Therefore, the overall resolution was determined from the Fourier shell correlation (FSC) of the map against the final refined atomic model (see below).

### Mycobacterium tuberculosis FtsZ

A schematic processing pipeline can be found in Supplementary Figure S6. Datasets were treated independently until the merging point indicated. Motion correction of micrograph movies was performed using Relion’s own implementation, motion corrected micrographs were then imported into cryoSPARC. Patch-based CTF estimation was performed (needed for the tilted images), a total of ∼5 M filament segments were picked using Topaz 0.2.4 (Bepler et al. 2019) via the cryoSPARC interface. Picks were extracted into 224^2^ pixel boxes, at 1.45 Å/pixel. 2D classification was used to remove bad particles, 3.9 M particles remaining were merged and used for non-uniform refinement to give a low-resolution map. Particles were recentred and re-extracted before another round of non-uniform refinement with little improvement. Further cleaning was carried out by reimporting the refinement results into Relion and performing 2D classification without alignment. 1.9 M particles remaining were reimported to cryoSPARC for a final non-uniform refinement to give the final medium-resolution map.

### Prosthecobacter dejongeii BtubA*B

#### GMPCPP-bound sample

A schematic processing pipeline can be found in Supplementary Figure S7. Tilted and untilted datasets were treated independently until the merging point indicated. Motion correction of micrograph movies was performed using Relion’s implementation, enabling Bayesian polishing at later steps. Per-patch CTF estimates, to account for sample tilt, were produced using Warp 1.0.9 (Tegunov and Cramer 2019). A total of 1.6 M filament segments (with Relion-compatible helical metadata) were picked at 45 Å intervals using Cryolo 1.7.4 in filament mode (Wagner et al. 2019), before extraction in 440^2^ Å^2^ boxes, binned to 2.2 Å/pixel. Two rounds of 2D classifications were performed before re-centring and re-extracting the remaining 1 M particles, unbinned (1.1 Å/pixel). These particle images and associated metadata were imported into cryoSPARC using pyem (Asarnow, Palovcak, and Cheng 2019). Homogenous refinement yielded a map with a reported resolution of 2.9 Å, however this was clearly a “mixed register” map – with both BtubA and BtubB subunits contributing to the density at all positions. The result of this mixed reconstruction (refined angles and shifts) was reimported into Relion. At this point alternate picks along filaments were removed (possible because helical metadata was retained), as if the initial particle segments had been picked and extracted at 90 Å spacing. 3D classification without aligment was performed on the remaining 500k particles (using the angles and shifts from the cryoSPARC refinement), and was able to separate the two registers present in the consensus map. 290k particles from classes corresponding to the two registers possibilities were re-centred and re-extracted with new centres shifted along the z-axis by half a monomer in either direction, before re-merging – yielding a set of images, angles and shifts corresponding to a reconstruction with a common register. This data was reimported to cryoSPARC for homogenous refinement, yielding a 3.2 Å map with clearly distinguishable BtubA and BtubB subunits, indicating success of the approach. This reconstruction was again imported into Relion for Bayesian polishing, before reimporting to cryoSPARC for homogenous refinement (2.7 Å) and, finally, non-uniform refinement (Punjani, Zhang, and Fleet 2020) producing the final 2.6 Å reconstruction.

#### GTP-bound sample

A schematic processing pipeline can be found in Supplementary Figure S9. Motion correction, CTF estimation and helical picking were all carried out using the Relion implementations. 750k particles were extracted in 440^2^ Å^2^ boxes, binned to 2.2 Å/pixel, before 2D classification. 460k remaining particles were imported to cryoSPARC for homogenous refinement to yield a low-resolution reconstruction (∼8 Å).

### Model building and refinement

Data and model statistics are summarised in Supplementary Table S1.

### *Drosophila melanogaster* αβ-tubulin heterodimers

GTP form: For model building, a *D. melanogaster* α/β-tubulin heterodimer was homology modelled using SWISSMODEL (Waterhouse et al. 2018). The dimer was placed manually in the cryo-EM map and adjusted manually in MAIN (Turk 2013), including the observed guanosine nucleotides. After manual building, the model was refined computationally using phenix.real_space_refine (Afonine et al. 2018). After several cycles of manual building and real-space-refinement, a satisfactory fit of the model to the map could be obtained and the model showed very good statistics: Molprobity score = 1.80, 85th percentile, with no Ramachandran outliers (V. B. Chen et al. 2010). GDP form: model building and refinement proceeded as for the α/β-tubulin heterodimer in the GTP form. Molprobity score = 1.39, 97th percentile, with no Ramachandran outliers.

### Drosophila melanogaster microtubule

Model building started with a PDB 3J1T *αβ-*tubulin heterodimer placed into a section of the final map, which was manually adjusted in MAIN (Turk 2013) and refined computationally using phenix.real_space_refine (Afonine et al. 2018). Once a satisfactory fit of the model to the map could be obtained, the heterodimer was repeatedly copied and placed in the map to describe three dimers in each of the 13 protofilaments. Finally, the entire model was refined using phenix.read_space_refine against the entire MT map. The model vs map FSC was 0.5 at ∼ 3.8 Å resolution and the model showed excellent statistics: Molprobity score = 1.74, 88^th^ percentile, with no Ramachandran outliers (V. B. Chen et al. 2010).

### Prosthecobacter dejongeii BtubA*B

For model building, a BtubAB heterodimer (PDB 2BTQ) was placed manually in the central section of the protofilament cryo-EM map and adjusted manually in MAIN (Turk 2013), including the observed GMPCPP nucleotide. The resolution of the map made it possible and indeed straightforward to determine the register of the protofilament with respect to BtubA and BtubB subunits. After manual building, the model was refined computationally using phenix.real_space_refine (Afonine et al. 2018). After several cycles of manual building and real-space-refinement, a satisfactory fit of the model to the map could be obtained. Three BtubAB dimers were then placed in the protofilament cryo-EM map and refined computationally using phenix.real_space_refine against the entire map. The model showed very good statistics: Molprobity score = 2.03, 74th percentile, with no Ramachandran outliers.

### Structural analysis

All analyses were carried out using custom scripts written in R, making extensive use of the bio3d (Grant et al. 2006) and tidyverse (Wickham 2017) packages.

Structure datasets were collected programmatically from the PDB (April 2021) by running individual phmmer (Eddy 2011) searches of PDB sequences with a representative from each of the subfamilies investigated.

Sequences from the downloaded PDB depositions were extracted and aligned as follows. Firstly, a high-quality representative structure was selected for each subfamily, and family members were aligned for each superfamily via a hybrid approach combining structural and sequence alignments, including information from large alignments of homologues, using the PROMALS3D web server with defaults (Pei, Kim, and Grishin 2008). Sequences were then aligned within each subfamily to the representative, using MUSCLE (Edgar 2004) (with some manual adjustments), before all sequences were combined into a super alignment on the basis of the representative alignment (with some manual adjustments).

Structures were annotated by downloading the Uniprot entry listed in the PDB annotation. Polymerisation state was assigned semi-automatically on the basis of experimental technique, but checked manually. Similarly, nucleotide state was assigned semi-automatically on the basis of ligand annotation in the PDB.

High-quality representatives of extant conformational states were selected by structurally aligning and clustering identically annotated (i.e. same sequence, same ligand, same polymerisation state) structures with RMSD < 1.0 Å, before choosing the highest resolution example from the cluster. Sometimes the cluster representative was manually adjusted to include a structure with a higher proportion of built residues, or rejected if other structure quality metrics were not very good.

Among the representatives, and using only the un-gapped positions in the super alignment of representatives within each superfamily, the structurally conserved core was found using the bio3d::core.find routine implementing Gerstein’s algorithm (Gerstein and Altman 1995; Gerstein and Chothia 1991; Grant et al. 2006). The structures were aligned on the core using bio3d::fit.xyz. Structure-based PCA was performed with bio3d::pca.pdbs.

## Supplementary Information

### Tables

**Supplementary Table S1.** Cryo-EM data collection and model statistics.

| Sample | Tubulin heterodimer<br>GDP | Tubulin heterodimer<br>GTP | Microtubule<br>GTP<br>13 protofilaments | BtubAB protofilament<br>GMPCPP<br>M-loop mutant:<br>A(R284G,K286D,<br>F287G) | FtsZ protofilament<br>GMPCPP |
| --- | --- | --- | --- | --- | --- |
| Organism | *D. melanogaster* | *D. melanogaster* | *D. melanogaster* | *P. dejongei* | *M. tuberculosis* |
| EM Data Bank map ID | EMD-14147 | EMD-14148 | EMD-14150 | EMD-14151 | Not deposited |
| PDB model ID | PDB 7QUC | PDB 7QUD | PDB 7QUP | PDB 7QUQ | Not deposited |
| **Data collection and processing** |  |  |  |  |  |
| Magnification | 64,000 | 64,000 | 81,000 | 81,000 | 105,000/81,000 |
| Voltage (kV) | 300 | 300 | 300 | 300 | 300 |
| Electron fluence (e-/Å<sup>2</sup>) | 40 | 40 | 35 | 44 | 40 |
| Defocus range (µm) | -1 to -3 (nominal) | -1 to -3 (nominal) | -1 to -3 (nominal) | -1 to -3 (nominal) | -1 to -3 (nominal) |
| Pixel size (Å) | 1.244 | 1.244 | 1.080 | 1.080 (some 40° tilted) | 0.86/1.47(40° tilted) |
| Symmetry imposed | C1 | C1 | local 13 pf MT | C1 | C1 |
| Initial particle images (no.) | 6,500,000 | 1,695,000 | 354,399 | 1,600,000 | 5,000,000 |
| Final particle images (no.) | 68,851 | 34,175 | 39,594 | 290,000 | 1,900,000 |
| Map resolution (Å) | 3.2 | 3.5 | Not gold standard | 2.6 (anisotropic) | ~7 (anisotropic) |
| FSC threshold | 0.143 | 0.143 | Not gold standard | 0.143 | 0.143 |
| **Model** |  |  |  |  |  |
| Initial model used<br>(PDB code) | SWISSMODEL | SWISSMODEL | 3J1T | 2BTQ | 6YM1 / 3VOA |
| Model resolution (Å) | 3.4 | 3.6 | 3.8 | 3.5 | Rigid body only |
| FSC threshold | 0.5 | 0.5 | 0.5 | 0.5 |  |
| Map sharpening B factor (Å<sup>2</sup>) | -81 | -61 | -175 | -33 |  |
| Model composition |  |  |  |  |  |
| Non-hydrogen atoms | 6,680 | 6,696 | 263,094 | 19,095 |  |
| Protein residues | 842 | 844 | 33,189 | 2,463 |  |
| Nucleic acid residues | 0 | 0 | 0 | 0 |  |
| Ligands | GTP: 1<br>GDP: 1<br>Mg: 1 | GTP: 2<br>Mg: 1 | GTP: 39<br>GDP: 39<br>Mg: 39 | G2P (GMPCPP): 6 |  |
| R.M.S. deviations |  |  |  |  |  |
| Bond lengths (Å) | 0.004 | 0.004 | 0.005 | 0.003 |  |
| Bond angles (°) | 0.557 | 0.610 | 0.744 | 0.754 |  |
| **Validation** |  |  |  |  |  |
| MolProbity score | 1.39 | 1.80 | 1.74 | 2.03 |  |
| Clashscore | 7.16 | 11.47 | 9.15 | 15.88 |  |
| Poor rotamers (%) | 0 | 2 | 2 | 3 |  |
| Ramachandran plot |  |  |  |  |  |
| Favored (%) | 98.2 | 96.5 | 96.2 | 95.3 |  |
| Allowed (%) | 1.8 | 3.5 | 3.8 | 4.5 |  |
| Disallowed (%) | 0 | 0 | 0 | 0 |  |

## Supplementary text

### Supplementary Text S1

**Notes on PCA**.

#### Details of structural survey

PC values and annotations for all representative structures are contained in Supplementary Files: supp_table_pca_act.csv and supp_table_pca_tub.csv.

|  | **Actins** | **Tubulins** |
| --- | --- | --- |
| Representatives used to search PDB | 5AEY_C, 5JLF_D, 5LJV_D, 5MW1_D, 4CZJ_B, 4A2B_A, 6F95_B, 4XHN_A | 1W59_A, 3CB2_A, 3ZID_A, 4EI7_A, 4XCQ_A, 3R4V_A, 2BTQ_A, 2BTQ_B, 3M89_A, 3VOA_A, 1RLU_A, 4B45_A, 5M8G_A, 5M8G_C |
| N of chains downloaded from PDB | 459 | 1441 |
| N of entries downloaded from PDB | 233 | 329 |
| N of chains used as cluster representatives for PCA | 107 | 165 |
| N of aligned, un-gapped “core” positions | 143 | 205 |
| N of invariant residues | 18 | 17 |

#### Commentary on selected actin structures

In this section and the one for tubulins, outliers in the PC subspaces are briefly described, along with all structures whose polymerisation state was annotated “special/ambiguous” (coloured green in Figures 2e and 5c, marked here with an asterisk “*”), often also outliers.

##### Outliers

2ZWH_A – Model for the eukaryotic F-actin structure (Oda et al. 2009), derived from fibre diffraction data; structure performs poorly on validation metrics and was refined without modern tools.

5YU8_A – Cofilin(severing protein)-decorated eukaryotic actin filament (Tanaka et al. 2018), i.e. highly unstable. Filament structure, but conformation is closer to the monomer cluster.

1HLU_A – Interesting eukaryotic actin open monomer, but structure performs poorly on validation metrics and was refined without modern tools.

5WFN_A – Eukaryotic actin complex with leiomodin, a nucleator protein that binds at multiple sites on the monomer. Interestingly the conformation is *more* open than most monomers (the opposite of e.g. 4A62_B).

4A62_B* – ParM complex with fragment of ParR, its filament nucleator. The monomeric structure adopts a conformation similar to the polymerised protein.

#### Commentary on selected tubulin structures

##### Outliers

5H5I_A* – FtsZ R29A point mutant in which the energy barrier between the subunit switch states has been removed (Fujita et al. 2017).

6UMK_A, 6UNX_A, 6LL5_A – three representatives of the somewhat unusual FtsZs from *Klebsiella pneumoniae* and *Escherichia coli*.

6BBN_D – aβ-tubulin from a curved tubulin complex induced by the kinesin-13 Kif2A.

1W59_A*, 1W59_B* – two FtsZ chains, identical in sequence, form a pseudo-polymerised dimer inside a crystal.

6B0C_A*, 6B0C_B* – these two chains form a polymer of tubulin heterodimers around the outside of a microtubule, linked via kinesin-13s.

6V5V_G* – structure of gamma-tubulin in the native human gamma-tubulin ring complex.

7ANZ_A*, 7ANZ_B* – two gamma tubulins within a structure of gamma-Tubulin Small Complex (gTuSC).

Four unusual α-tubulin conformations, from polymer structures but which are found in the monomeric conformation cluster:

1TVK_A – electron crystallography structure.

6REV_A – Cryo-EM structure of human doublecortin (MT stabiliser) bound to 13-pf GDP-MT.

1JFF_A – electron crystallography structure of zinc-induced sheets.

6CVJ_A – Tau bound to MT by cryo-EM. Possibly a refinement mistake.

And the reverse, two α-tubulin monomer structures that lie close to the filament cluster.

6GWC_A – from a heterodimer in complex with a synthetic protein binding partner.

FFB_A – from a heterodimer in complex with a TOG MT depolymerase domain.

### Supplementary Text S2

**Proteins used in this study**.

#### *Drosophila melanogaster* tubulins

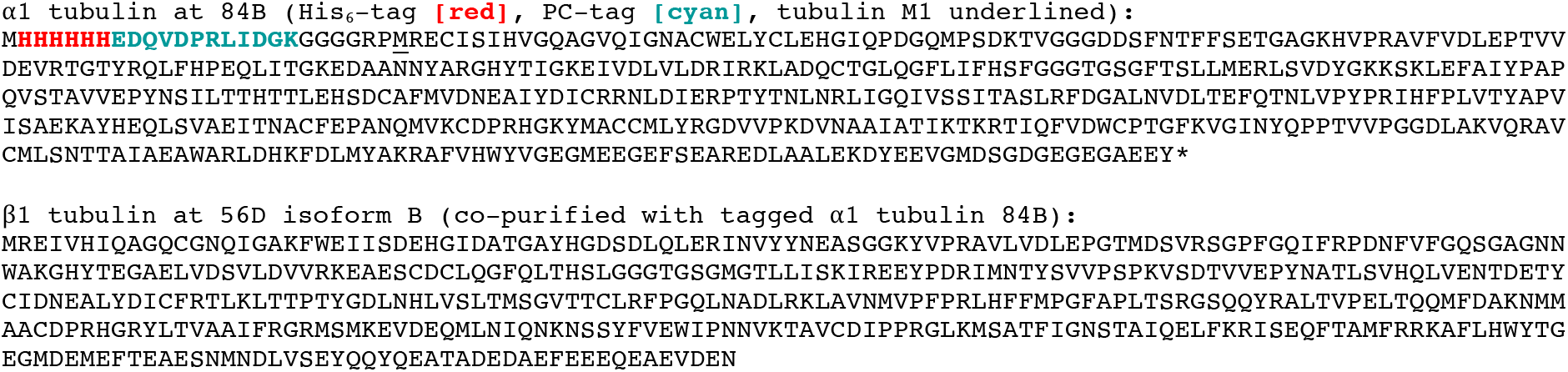

#### *Mycobacterium tuberculosis* FtsZ

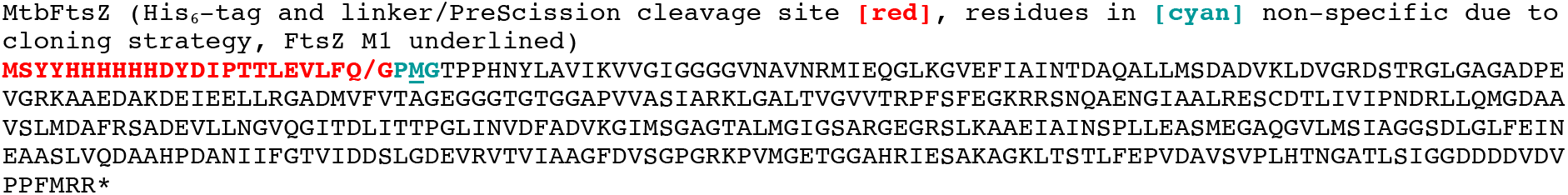

#### *Prosthecobacter dejongeii* BtubA*B

##### BtubA*, (R284G, R286D, F287G, M-loop mutations underlined)

MKVNNTIVVSIGQAGNQIAASFWKTVCLEHGIDPLTGQTAPGVAPRGNWSSFFSKLGESSSGSYVPRAIMVDLEPSVIDNVKATSGSLFNPANLISRTEG AGGNFAVGYLGAGREVLPEVMSRLDYEIDKCDNVGGIIVLHAIGGGTGSGFGALLIESLKEKYGEIPVLSCAVLPSPQVSSVVTEPYNTVFALNTLRRSA DACLIFDNEALFDLAHRKWNIESPTVDDLNLLITEALAGITASMRFSGFLTVEISLRELLTNLVPQPSLHFLMCAFAPLTPPD<u>G</u>S<u>DG</u>EELGIEEMIKSLF DNGSVFAACSPMEGRFLSTAVLYRGIMEDKPLADAALAAMREKLPLTYWIPTAFKIGYVEQPGISHRKSMVLLANNTEIARVLDRICHNFDKLWQRKAFANWYLNEGMSEEQINVLRASAQELVQSYQVAEESGAKAKVQDSAGDTGMRAAAAGVSDDARGSMSLRDLVDRRR

##### BtubB

MREILSIHVGQCGNQIADSFWRLALREHGLTEAGTLKEGSNAAANSNMEVFFHKVRDGKYVPRAVLVDLEPGVIARIEGGDMSQLFDESSIVRKIPGAANNWARGYNVEGEKVIDQIMNVIDSAVEKTKGLQGFLMTHSIGGGSGSGLGSLILERLRQAYPKKRIFTFSVVPSPLISDSAVEPYNAILTLQRILDNADGAVLLDNEALFRIAKAKLNRSPNYMDLNNIIALIVSSVTASLRFPGKLNTDLSEFVTNLVPFPGNHFLTASFAPMRGAGQEGQVRTNFPDLARETFAQDNFTAAIDWQQGVYLAASALFRGDVKAKDVDENMATIRKSLNYASYMPASGGLKLGYAETAPEGFASSGLALVNHTGIAAVFERLIAQFDIMFDNHAYTHWYENAGVSRDMMAKARNQIATLAQSYRDAS

## Supplementary figures

**Supplementary Figure S1.**
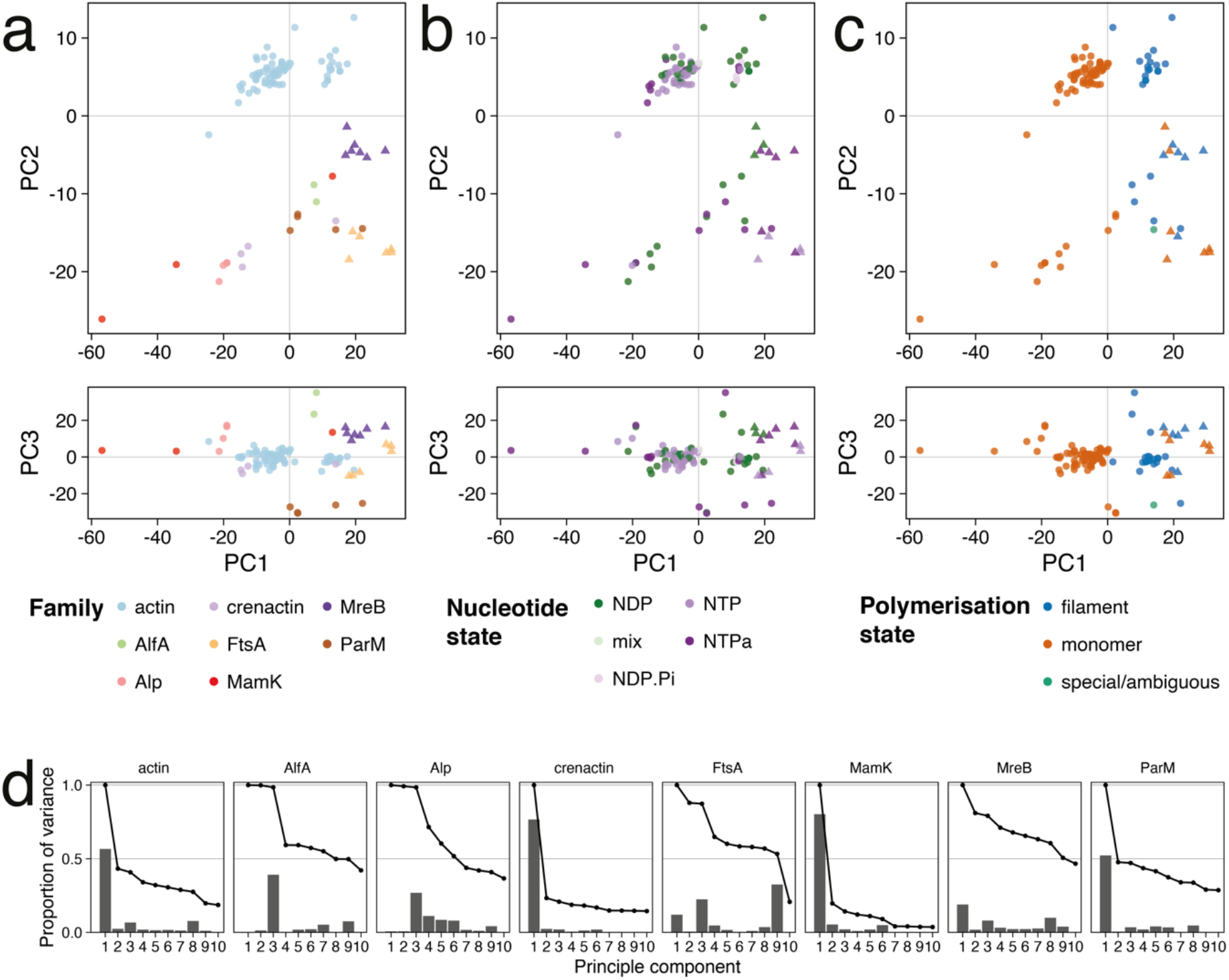
Actin PCA supplements. (a-c) Actin PC subspaces as in Figure 2, with the PC1-PC3 subspaces added below each plot. (d) Proportion of variance within each family’s Cαs explained by each of the superfamily PCs. PC1 is descriptive for the (cytomotive) families with representatives of both polymerised and un-polymerised subunits. PC2 mostly describes differences between families so is not descriptive for any given family.

**Supplementary Figure S2.**
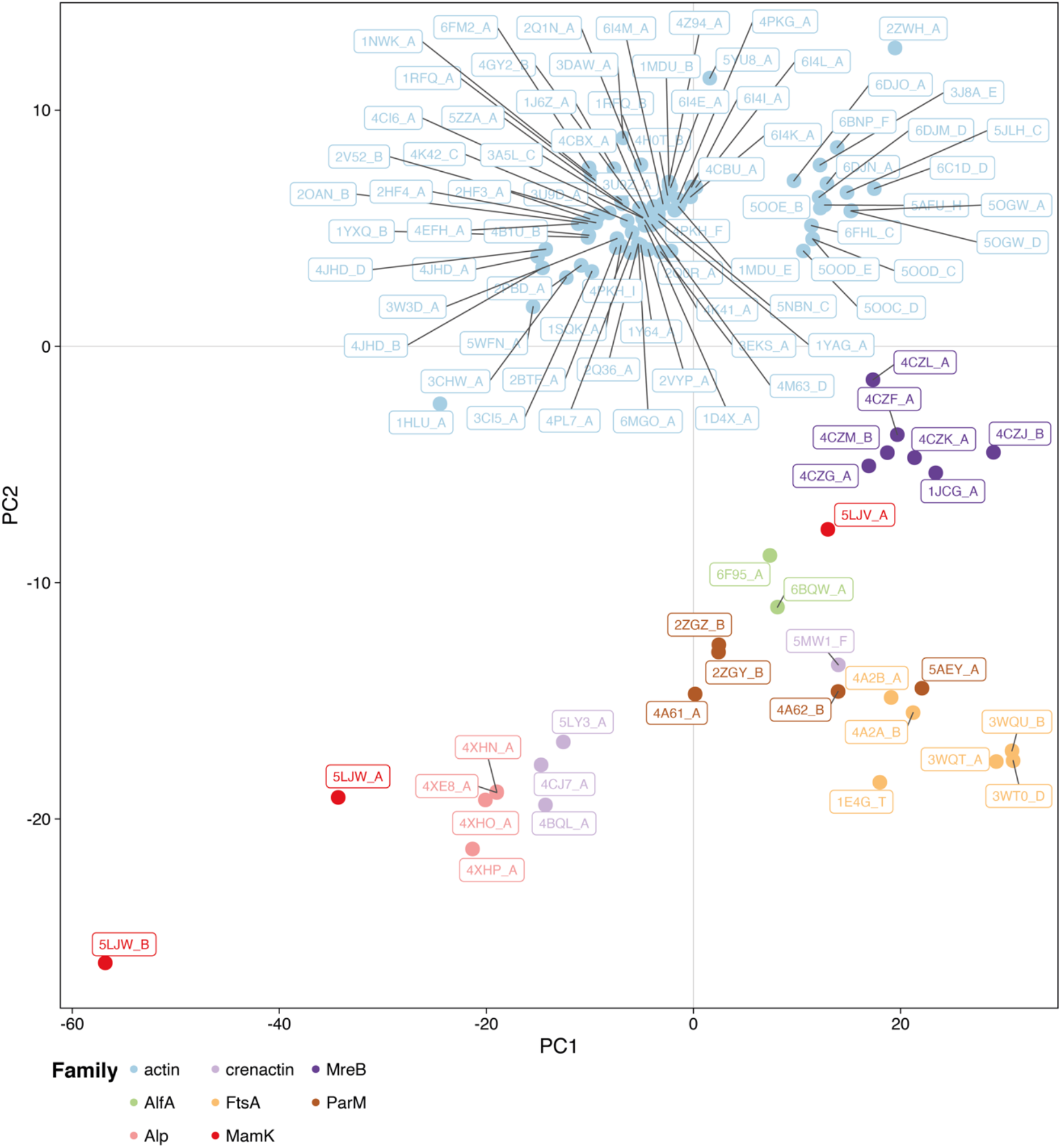
Actin PCA with labelled structures. Actin PC1-PC2 subspace as in Figure 2c, with each representative structure labelled in the format ‘“PDB accession”_”chain id”‘.

**Supplementary Figure S3.**
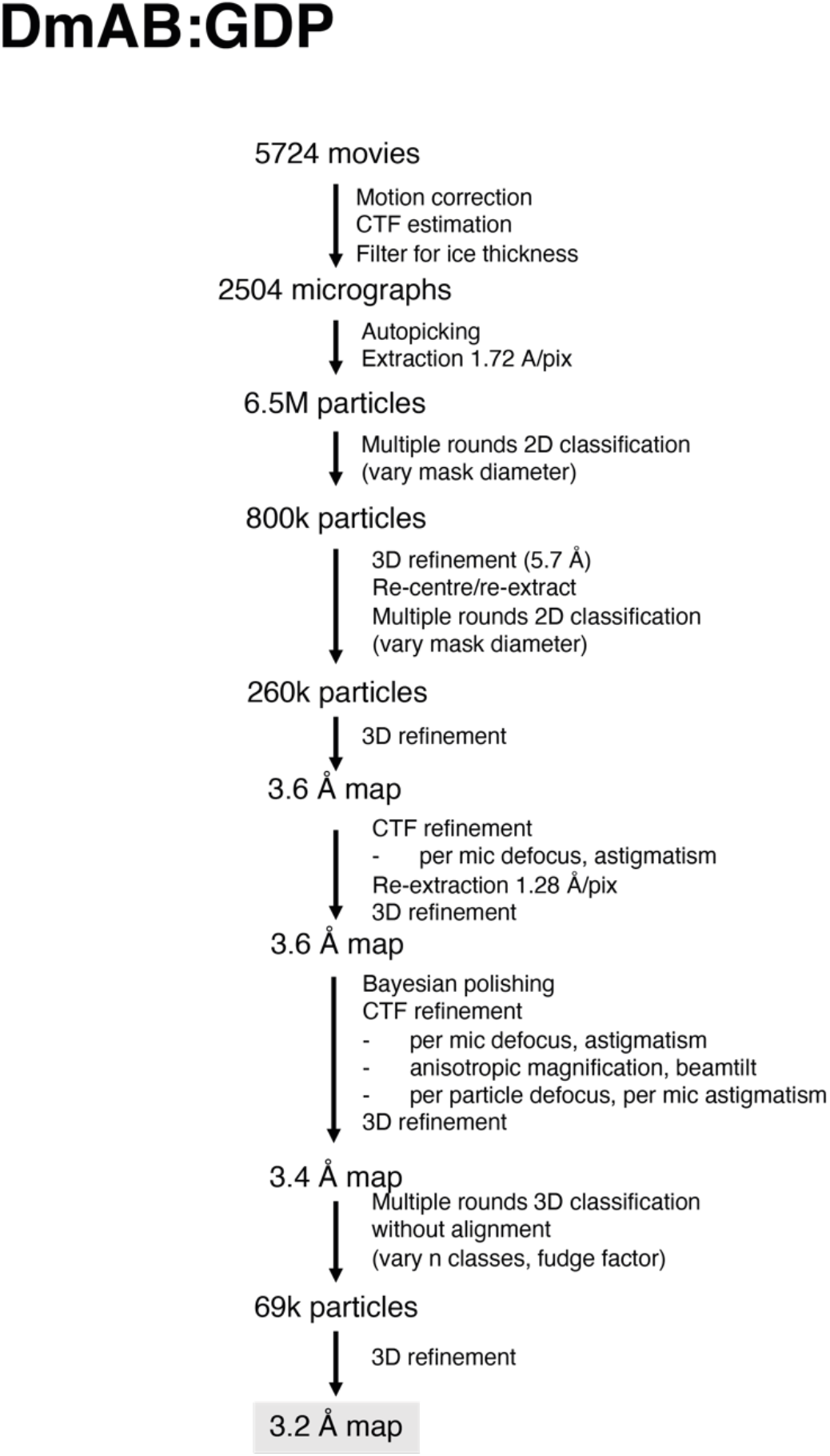
*Drosophila melanogaster* (*Dm*) tubulin GDP cryo-EM processing scheme.

**Supplementary Figure S4.**
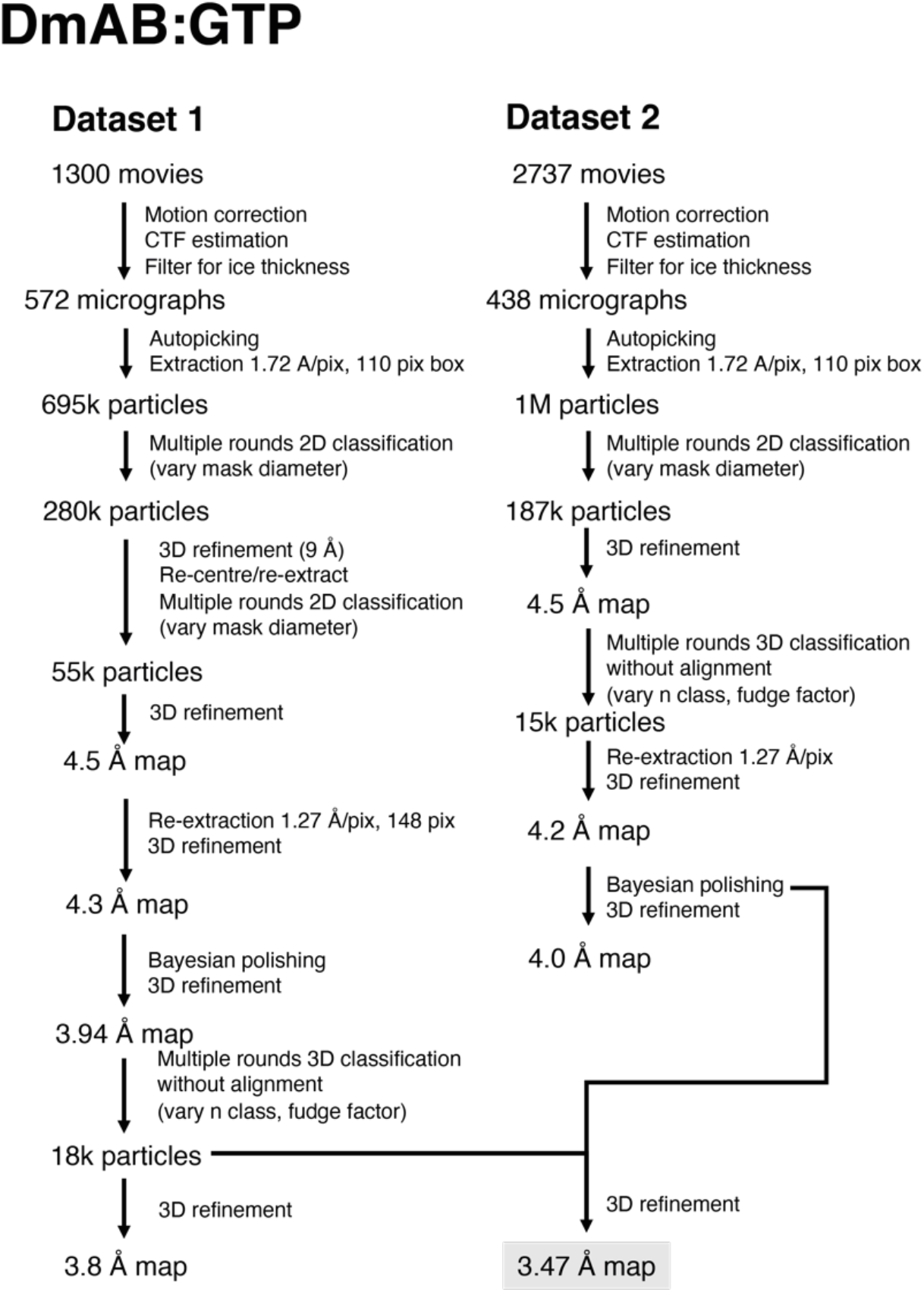
*Drosophila melanogaster* (*Dm*) tubulin GTP cryo-EM processing scheme.

**Supplementary Figure S5.**
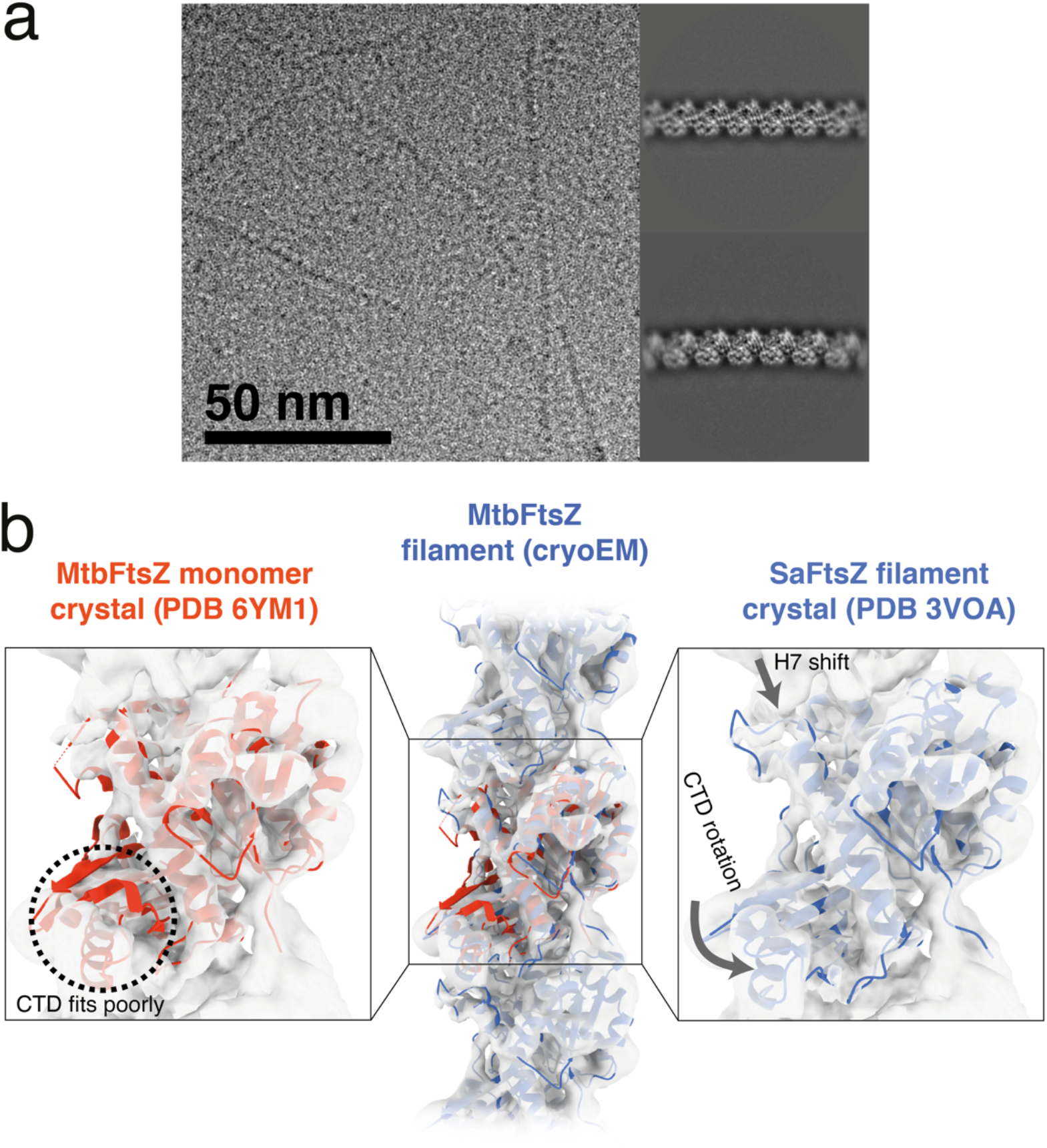
Cryo-EM structure of *Mycobacterium tuberculosis* (*Mtb*) FtsZ, a single protofilament tubulin, reveals a strict assembly switch. (a) Cryo-EM study of *Mycobacterium tuberculosis* (*Mtb*) FtsZ filaments. Representative micrograph and 2D class averages. Processing scheme can be found in Supplementary Figure S6. (b) Medium-resolution cryo-EM map of FtsZ filament from *M. tuberculosis*, reveals the assembly switch upon polymerisation. The map is compatible with the polymeric *S. aureus* FtsZ crystal structure (blue), but not the monomeric *M. tuberculosis* structure (orange). Map resolution is limited because of severe preferred orientation of the filaments after vitrification and the fact that the filaments are not helical (as was the case for *E. coli* FtsZ filaments, described in (J. M. Wagstaff et al. 2017))

**Supplementary Figure S6.**
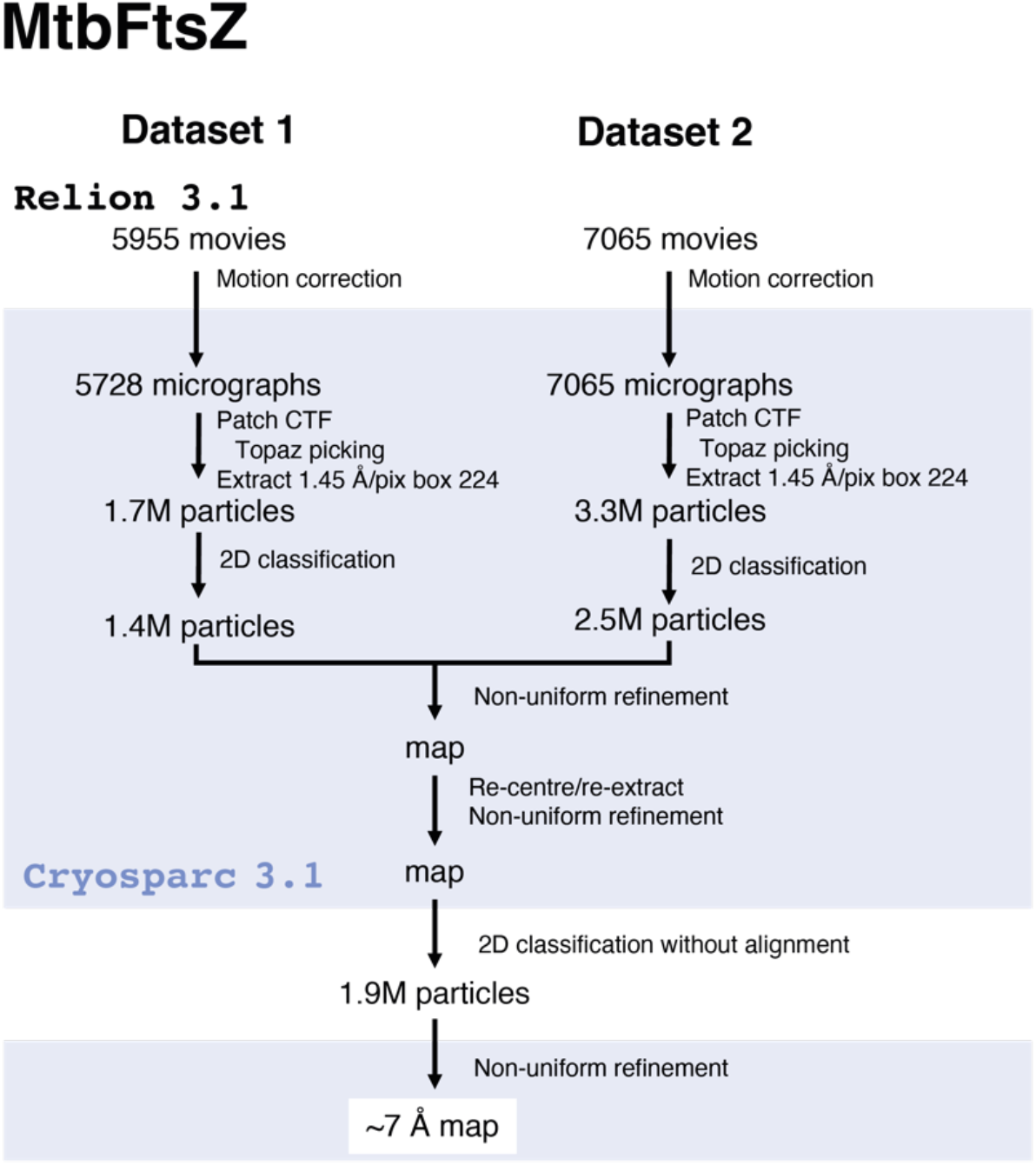
*Mycobacterium tuberculosis* (*Mtb*) FtsZ:GMPCPP cryo-EM processing scheme.

**Supplementary Figure S7.**
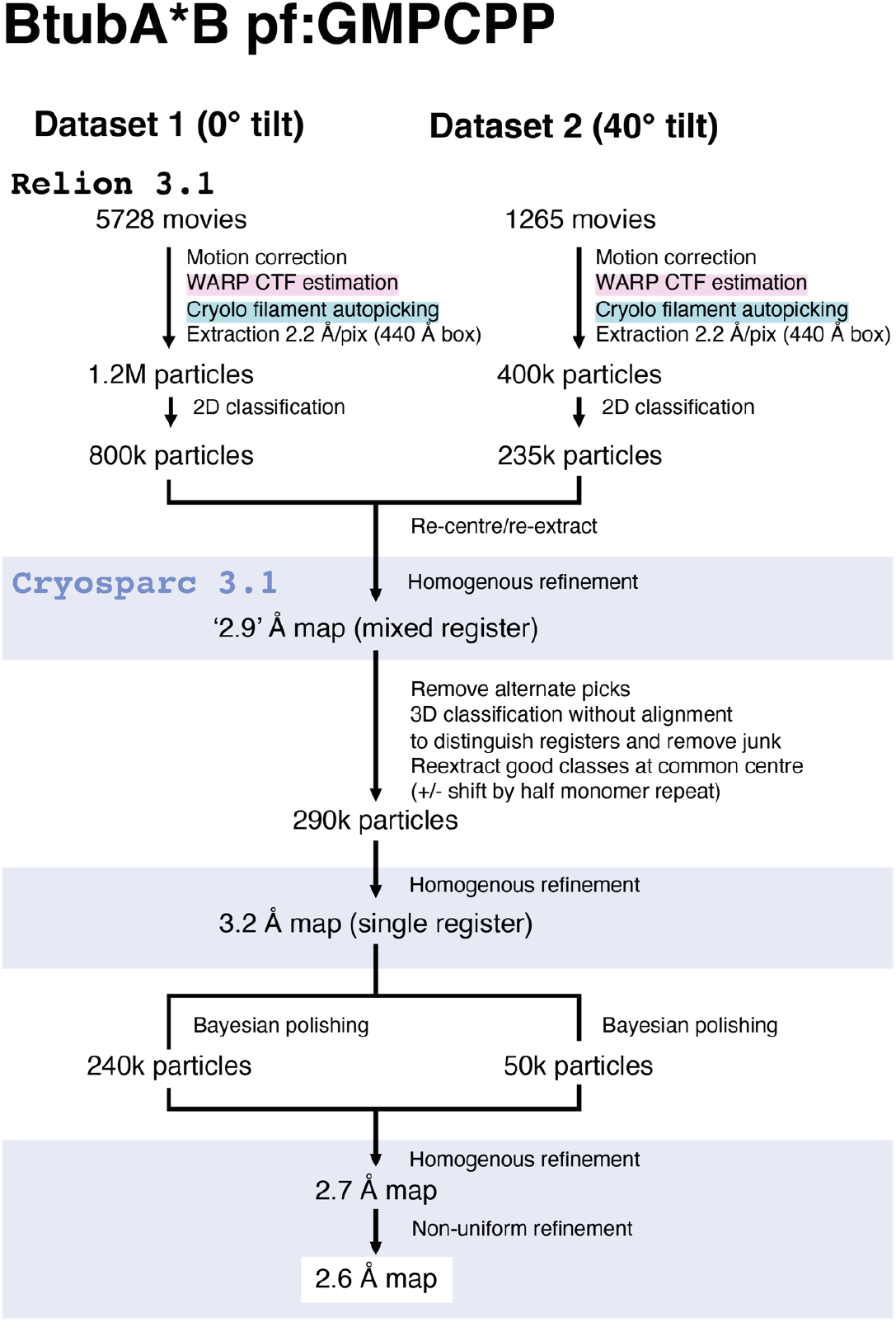
BtubA*B:GMPCPP cryo-EM processing scheme.

**Supplementary Figure S8.**
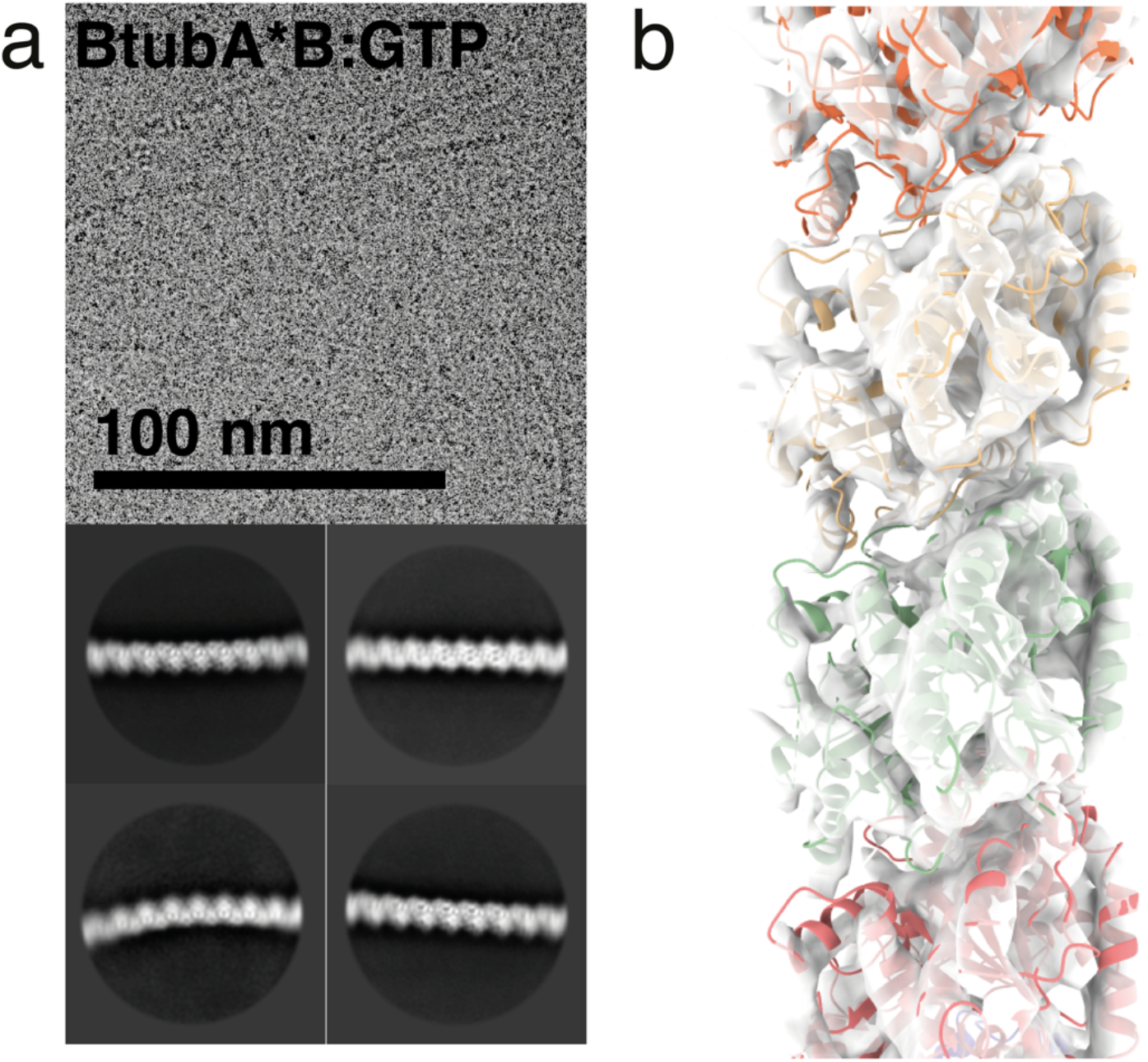
BtubA*B polymerises with GTP to form single-stranded filaments. (a) Cryo-EM study of short and curved BtubAB filaments assembled with GTP. Representative micrograph and 2D class averages. (b) Medium resolution cryo-EM map (∼ 8 Å resolution) of BtubA*B:GTP, with model of BtubAB filament rigid body fitted.

**Supplementary Figure S9.**
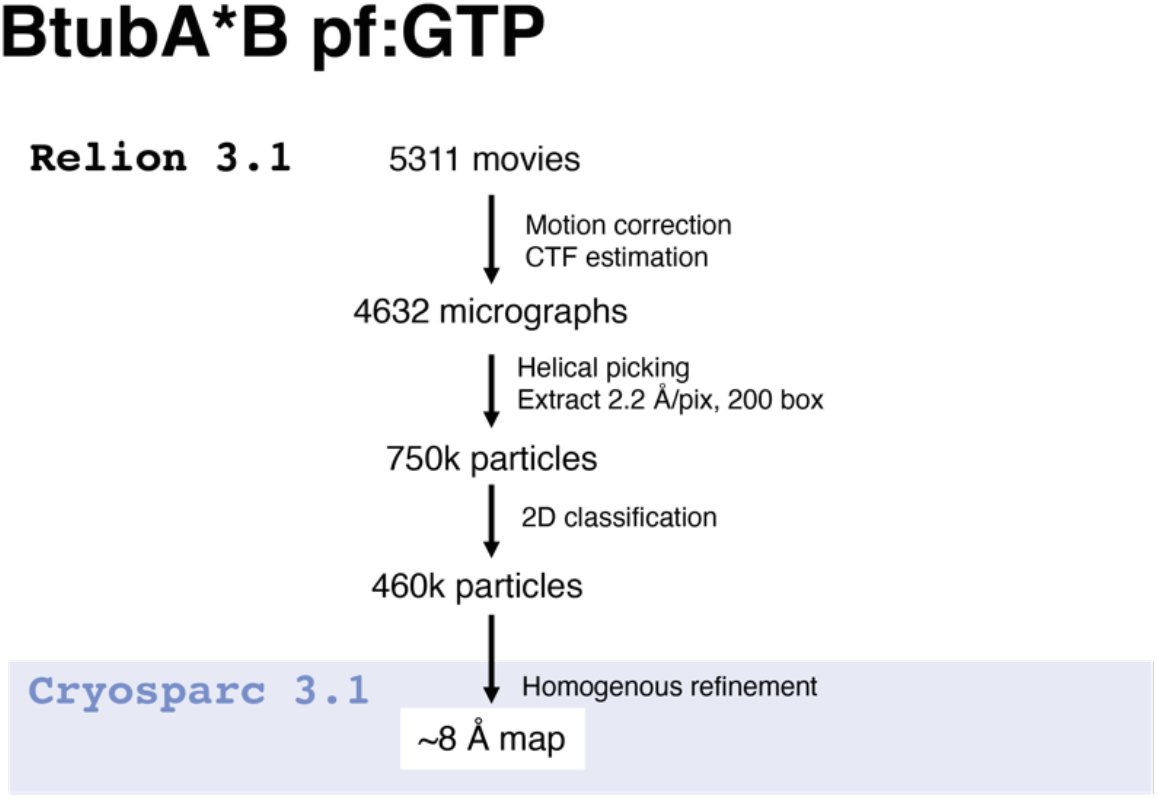
BtubA*B:GTP cryo-EM processing scheme.

**Supplementary Figure S10.**
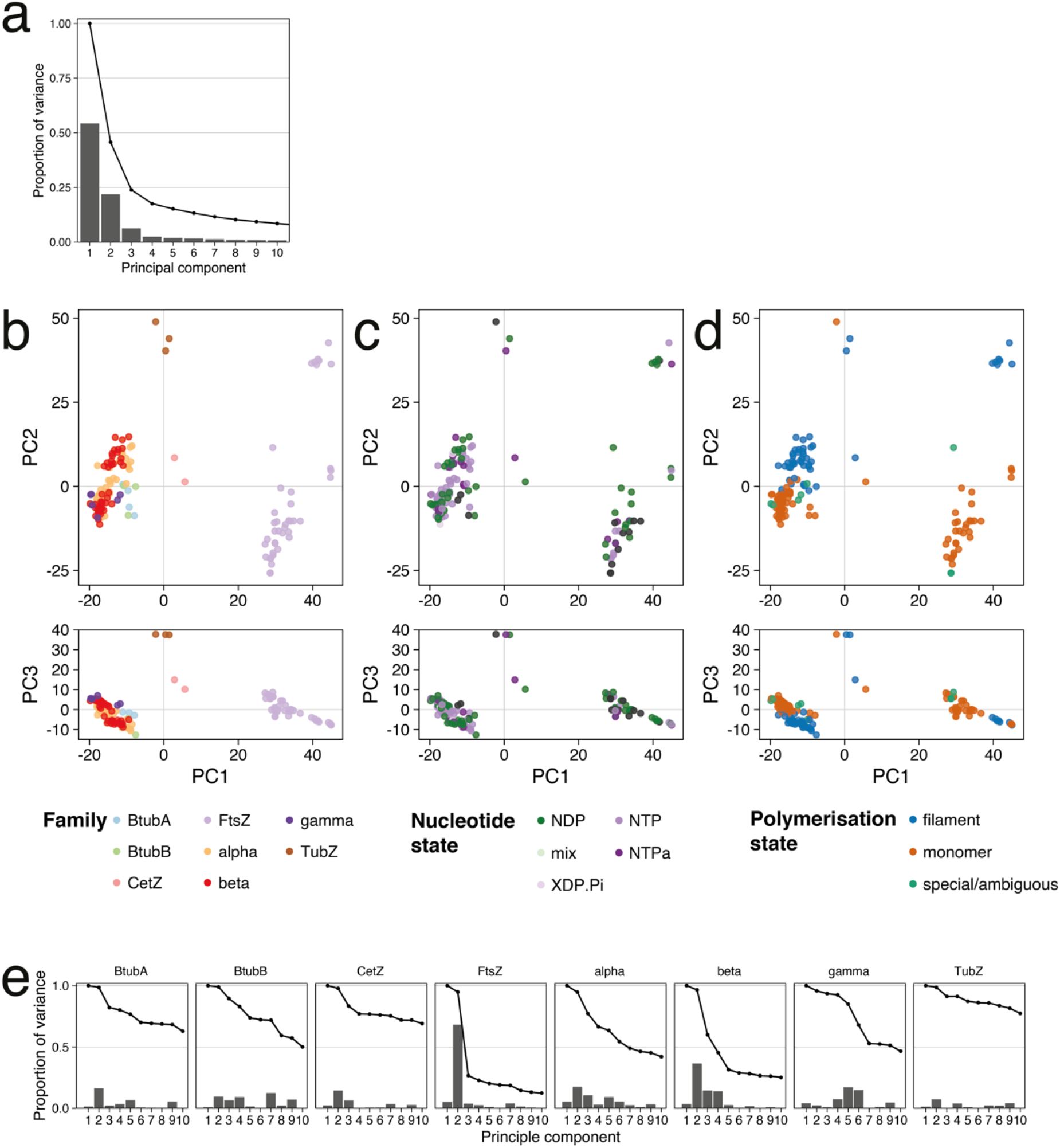
Tubulin PCA supplements. (a) Proportion of variance across all tubulin structures explained by the principal components (PC). (b-d) Tubulin PC subspaces as in Figure 5, with the PC1-PC3 subspaces added below each plot. (e) Proportion of variance within each family’s Cαs explained by each of the superfamily PCs. PC2 is descriptive for the (cytomotive) families with representatives of both polymerised and un-polymerised subunits because it describes the polymerisation switch. PC1 mostly describes differences between families so is not descriptive for any given family.

**Supplementary Figure S11.**
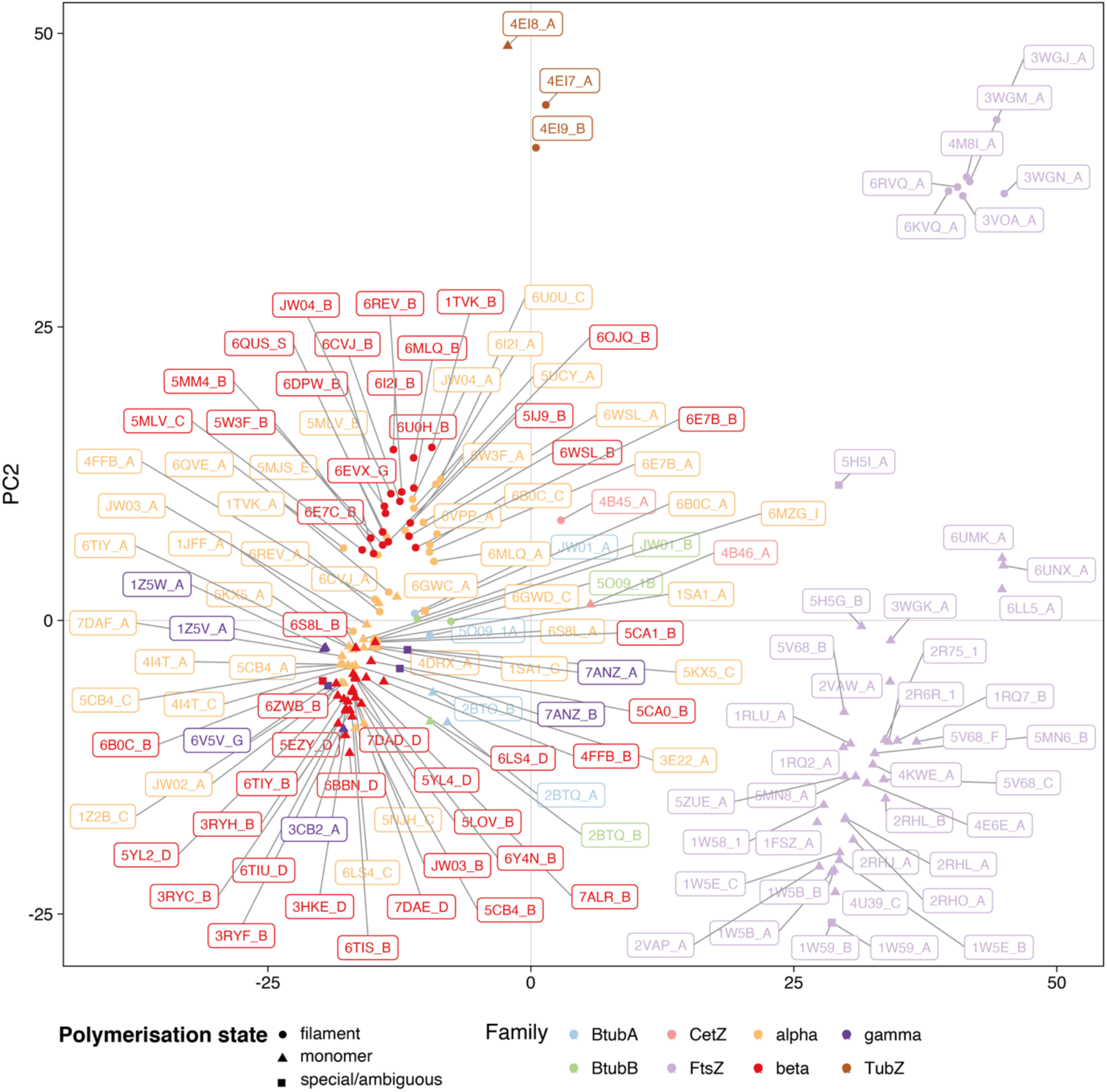
Tubulin PC1-PC2 subspace with labelled structures. Tubulin PC1-PC2 subspace as in Figure 5a, with each representative structure labelled in the format ‘“PDB accession”_”chain id”‘.

**Supplementary Figure S12.**
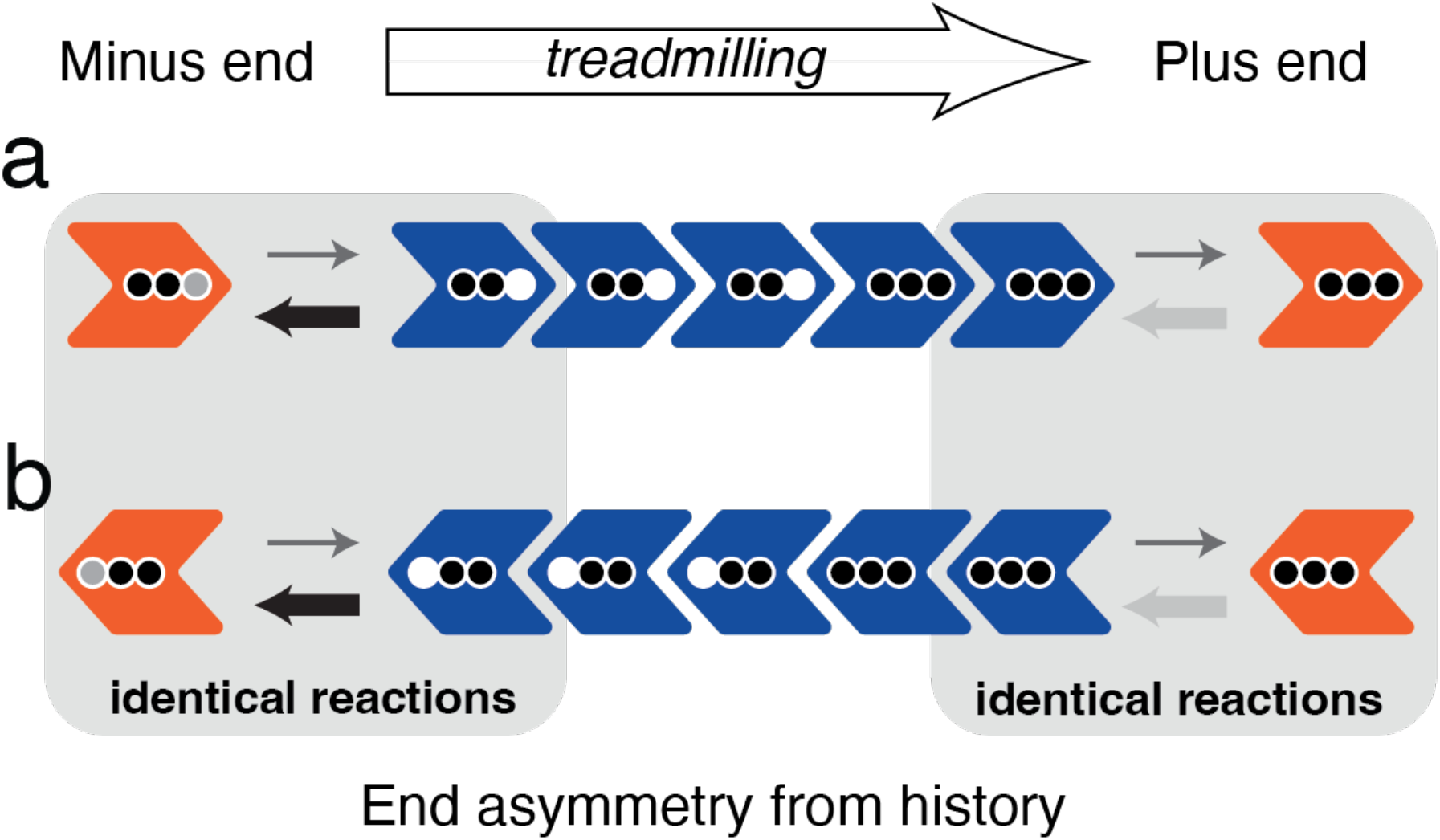
Un-coupled kinetic and structural polarities in a filament without a cytomotive switch. Two possible outcomes for dynamic polymerisation of one species of rigid subunits as in Fig 6a. These subunits can treadmill, but the direction in which they do so will be arbitrarily set by stochastically formed gradients of nucleotide hydrolysis within the filament. Thus, the kinetic (plus/minus) polarity will be uncoupled from the structural (pointed/notched end of chevrons) polarity. Such a filament will be less useful for many cytomotive tasks, as specific end binding is often used for positioning sub-cellular components.

